# The effect of background noise and its removal on the analysis of single-cell expression data

**DOI:** 10.1101/2022.11.16.516780

**Authors:** Philipp Janssen, Zane Kliesmete, Beate Vieth, Xian Adiconis, Sean Simmons, Jamie Marshall, Cristin McCabe, Holger Heyn, Joshua Z. Levin, Wolfgang Enard, Ines Hellmann

## Abstract

**BACKGROUND:** In droplet-based single-cell and single-nucleus RNA-seq experiments, not all reads associated with one cell barcode originate from the encapsulated cell. Such background noise is attributed to spillage from cell-free ambient RNA or barcode swapping events. Here, we characterize this background noise exemplified by three single-cell RNA-seq (scRNA-seq) and two single-nucleus RNA-seq (snRNA-seq) replicates of mouse kidney cells. For each experiment, kidney cells from two mouse subspecies were pooled, allowing to identify cross-genotype contaminating molecules and estimate the levels of background noise.

**RESULTS:** We find that background noise is highly variable across replicates and individual cells, making up on average 3-35% of the total counts (UMIs) per cell and show that this has a considerable impact on the specificity and detectability of marker genes. In search of the source of background noise, we find that expression profiles of cell-free droplets are very similar to expression profiles of cross-genotype contamination and hence that the majority of background molecules originates from ambient RNA. Finally, we use our genotype-based estimates to evaluate the performance of three methods (CellBender, DecontX, SoupX) that are designed to quantify and remove background noise. We find that CellBender provides the most precise estimates of background noise levels and also yields the highest improvement for marker gene detection. By contrast, clustering and classification of cells are fairly robust towards background noise and only small improvements can be achieved by background removal that may come at the cost of distortions in fine structure.

**CONCLUSION:** Our findings help to better understand the extent, sources and impact of background noise in single-cell experiments and provide guidance on how to deal with it.

## Background

Single cell and single nucleus RNA-seq (scRNA-seq, snRNA-seq) are in the process of revolutionizing medical and biological research. The typically sparse coverage per cell and gene is compensated by the capability of analyzing thousands of cells in one experiment. In droplet-based protocols such as 10x Chromium, this is achieved by encapsulating single cells in droplets together with beads that carry oligonucleotides. These usually consist of a oligo(dT) sequence which is used for priming reverse transcription, a bead-specific barcode that tags all transcripts encapsulated within the droplet and unique molecular identifiers (UMIs) that enable the removal of amplification noise [1, 2, 3]. As proof of principle that each droplet encapsulates only one cell, it is common to use mixtures of cells from human and mouse [3]. Thus doublets, droplets containing two cells, can be readily identified as they have an approximately even mixture of mouse and human transcripts. However, barcodes for which the clear majority of reads is either mouse or human, still contain a small fraction of reads from the other species [3, 4, 5]. Furthermore, presumably empty droplets also yield sequence reads [4].

One potential source of such contaminating reads or background noise is cell-free’ambient’ RNA that leaked from broken cells into the suspension. The other potential source are chimeric cDNA molecules that can arise during library preparation due to so-called’barcode swapping’. The pooling of barcode tagged cDNA after reverse transcription but before PCR amplification, is a decisive step to achieve high throughput. However, if amplification of tagged cDNA molecules occurs from unremoved oligonucleotides from other beads or from incompletely extended PCR products (originally called template jumping [6]), this generates a chimeric molecule with a “swapped” barcode and UMI [7, 8]. When sequencing this molecule, the cDNA is assigned to the wrong barcode and hence’contaminates’ the expression profile of a cell. Another type of barcode swapping can occur during PCR amplification on a patterned Illumina flowcell before sequencing [9] with the same effects, although double indexing of Illumina libraries has reduced this problem substantially. This said, here we focus on barcode swapping that occurs during library preparation.

Irrespective of the source of background noise, its presence can interfere with analyses. For starters, background noise reduces the separability of cell type clusters as well as the power to pinpoint important (marker) genes via differential expression analysis. Moreover, reads from cell type-specific marker genes spill over to cells of other types, thus yielding novel marker combinations and hence implying the presence of novel cell types [10, 8]. Besides, background noise can also confound differential expression analysis between samples, e.g. when looking for expression changes within a cell type between two conditions. Varying amounts of background noise or differences in the cell type composition between conditions can result in dissimilar background profiles, which might generate false positives when identifying differentially expressed genes. To alleviate such problems during downstream analysis, algorithms to estimate and correct for the amounts of background noise have been developed.

SoupX estimates the contamination fraction per cell using marker genes and then deconvolutes the expression profiles using empty droplets as an estimate of the background noise profile [11]. In contrast, DecontX defaults to model the fraction of background noise in a cell by fitting a mixture distribution based on the clusters of good cells [8], but also allows the user to provide a custom background profile, e.g. from empty droplets. CellBender requires the expression profiles measured in empty droplets to estimate the mean and variance of the background noise profile originating from ambient RNA. In addition, CellBender explicitly models the barcode swapping contribution using mixture profiles of the’good’ cells [4].

In order to evaluate method performance, one dataset of an even mix between one mouse and one human cell line [3] is commonly used to get an experimentally determined lower bound of background noise levels that is identified as counts covering genes from the other species [4, 8, 11, 12]. Since this dataset is lacking in cell type diversity, it is common to additionally evaluate performance based on other datasets that have a complex cell type mixture and where most cell types have well known profiles with exclusive marker genes. In such studies the performance test is whether the model removes the expression of the exclusive marker genes from the other cell types. In both cases, the feature space of the contamination does not overlap with the endogenous cell feature space. Mouse and human are too diverged, so that mouse reads only map to mouse genes and human reads only to human genes. Similarly, when using marker genes it is assumed that they are exclusively expressed in only one cell type, hence the features that are used for background inference are again not overlapping. However, in reality background noise will mostly induce shifts in expression levels that cannot be described in a binary on or off sense and it remains unclear how background correction will affect those profiles.

Here, we use a mouse kidney dataset representing a complex cell type mixture from three mouse strains of two subspecies, *Mus musculus domesticus* and *M*.*m*.*castaneus*. From both subspecies, inbred strains were used and thus we can distinguish exogenous and endogenous counts for the same features using known homozygous SNPs [13]. Hence, this dataset serves as a much more realistic experimental standard, providing a ground truth in a complex setting with multiple cell types which allows to analyze the variability, the source and the impact of background noise on single cell analysis. Moreover, this dataset enables us to better benchmark existing background removal methods.

### Mouse kidney single cell and single nucleus RNA-seq data

We obtained three replicates for single cell RNA-seq (rep1-3) data and two replicates for single nucleus RNA-seq (snRNA-seq, nuc2 & nuc3) data from the same samples that were used in scRNA-seq replicates 2 and 3, respectively. Each replicate consists of one channel of 10x [3] in which cells from dissociated kidneys of three mice each were pooled: one *M*.*m. castaneus* from the strain CAST/EiJ (CAST) and two *M*.*m. domesticus*, one from the strain C57BL/6J (BL6) and one from the strain 129S1/SvImJ (SvImJ) (Figure 1A). Based on known homozygous SNPs that distinguish subspecies and strains, we assigned cells to mice (Figure 1B). In total, we identified *>* 40, 000 informative SNPs of which the majority (32,000) separates the subspecies and ∼ 10, 000 SNPs distinguish the two *M*.*m. domesticus* strains (Figure 1C). On average, each cell had sufficient coverage for ∼ 1, 000 informative SNPs (∼ 20% of total UMIs per cell) to provide us with unambiguous genotype calls for those sites. The coverage for the nuc2 data was much lower with only ∼ 100 SNPs (Figure 1D).

**Figure 1.**
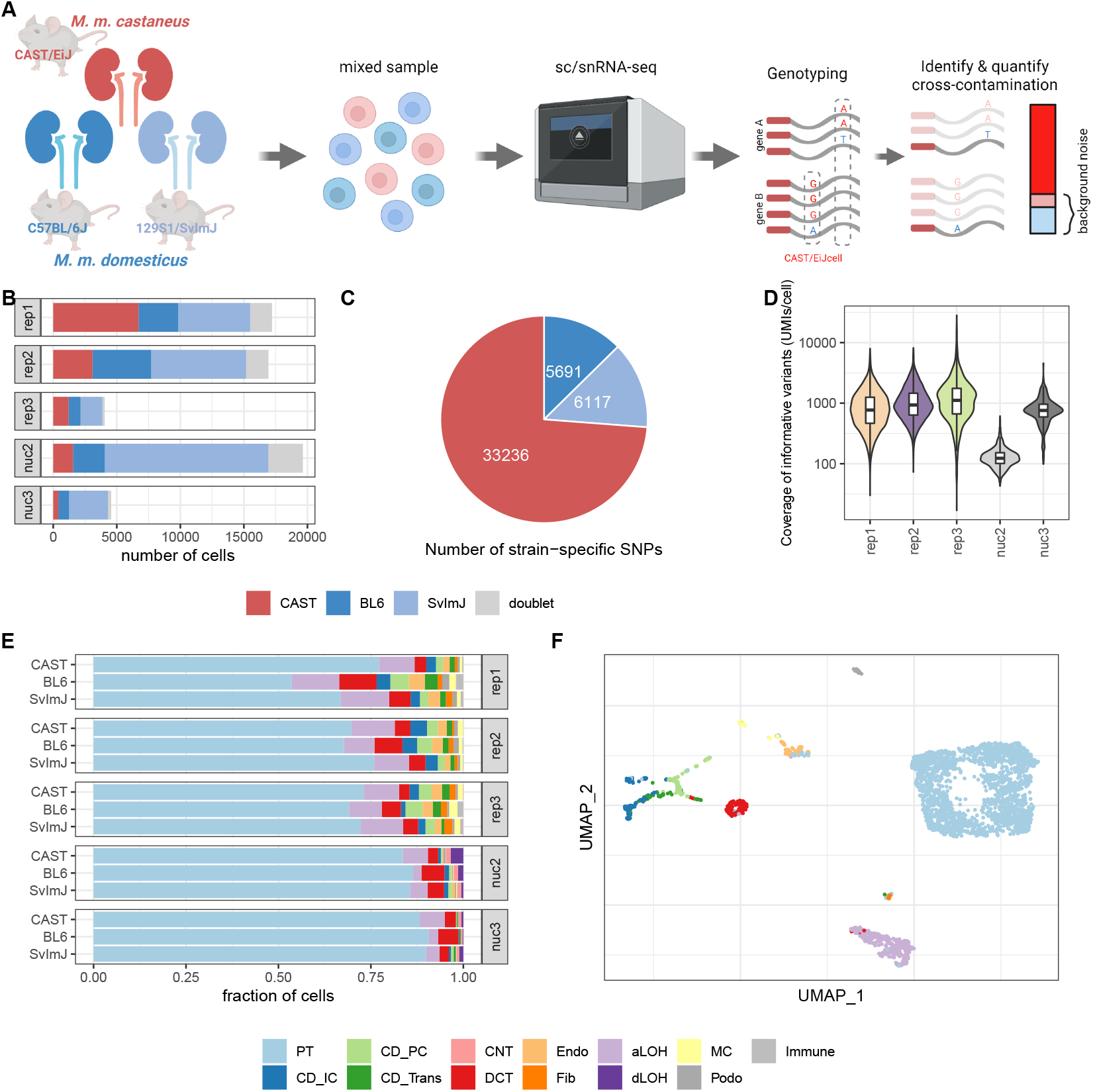
Generation of mouse strain mixture datasets to quantify background noise. A) Experimental design. B) Strain composition in 5 different replicates, subjected to scRNA-seq (rep1-3) or snRNA-seq (nuc2,nuc3). The replicates rep2 & nuc2 and rep3 & nuc3 were generated from the same samples each. CAST: CAST/EiJ strain; BL6: C57BL/6J strain; SvImJ: 129S1/SvImJ. C) Number of homozygous SNPs with a coverage of more than 100 UMIs that distinguish one strain from the other two. D) Per cell coverage in *M*.*m. castaneus* cells of informative variants that distinguish *M*.*m*.*castaneus* and *M*.*m*.*domesticus* E) Cell type composition per replicate and strain; labels were obtained by reference-based classification using mouse kidney data from Denisenko et al. [14] as reference. F) UMAP visualization of *M*.*m*.*castaneus* cells in single-cell replicate 2, colored by assigned cell type. PT: proximal tubule; CD_IC: intercalated cells of collecting duct; CD_PC: principal cells of collecting duct; CD_Trans: transitional cells of collecting duct; CNT: connecting tubule; DCT: distal convoluted tubule; Endo: endothelial; Fib: fibroblasts; aLOH: ascending loop of Henle; dLOH: descending loop of Henle; MC: mesangial cells; Podo: podocytes

Overall, each experiment yielded 5,000-20,000 good cells with 9-43% *M*.*m. castaneus* (Figure 1B). Thus, the majority of background noise in any *M*.*m. castaneus* cell is expected to be from *M*.*m. domesticus* and therefore we expect that genotype-based estimates of cellwise amounts of background noise for *M*.*m. castaneus* to be fairly accurate (Supplementary figure S1). Hence from here on out we focus on *M*.*m. castaneus* cells for the analysis of the origins of background noise and also as the ground truth for benchmarking background removal methods.

This dataset has two advantages over the commonly used mouse-human mix [3]. Firstly, the kidney data have a high cell type diversity. Using the data from Denisenko et al. [14] as reference dataset for kidney cell types, we could identify 13 cell types. Encouragingly, the cell type composition is very similar across mouse strains as well as replicates with proximal tubule cells constituting 66-89% of the cells (Figure 1E,F, Supplementary Figure S2). Secondly, due to the higher similarity of the mouse subspecies, we can identify contaminating reads for the same features. ∼ 7, 000 genes carry at least one informative SNP about the subspecies allowing us to quantify contaminating reads from the other mice.

### Background noise fractions differ between replicates and cells

Around 20% of the UMI counts are from molecules that contain a SNP that is informative about the subspecies of origin. We quantify in each cell how often an endogenous *M*.*m. castaneus* allele or a foreign *M*.*m. domesticus* allele was covered. Assuming that the count fractions covering the SNPs are representative of the whole cell, we detect a median of 2%-27% counts from the foreign genotype over all cells per experiment (Supplementary Figure S3A). This observed cross-genotype contamination fraction represents a lower bound of the overall amounts of background noise. As suggested in Heaton et al. [15], we then integrate over the foreign allele fractions of all informative SNPs to obtain a maximum likelihood estimate of the background noise fraction (*ρ*_*cell*_) of each cell that extrapolates to also include contamination from the same genotype (see Methods, Supplementary figure S1). Based on these estimates, we find that background noise levels vary considerably between replicates and do not appear to depend on the overall success of the experiment measured as the cell yield per lane (Figure 2). For example in scRNA-seq rep3 (3,900 cells), we detected overall the fewest good cells, but most of those cells had less than 3% background noise, while the much more successful rep2 (15,000 cells) we estimated the median background noise level at around 11% (Figure 2A). This said, the snRNA-seq data generated from frozen tissue have much higher background levels than the corresponding scRNA-seq replicates - 35% in nuc2 vs. 11% rep2 and 17% in nuc3 vs. 3% in rep3. The number of contaminating RNA-molecules (UMIs) depends only weakly on the sequencing depth of the cell (Figure 2B). Such a weak correlation could be explained by variation in the capture efficiency in each droplet. An alternative, but not mutually exclusive explanation of such a correlation could be that the source of some contaminating molecules is barcode swapping that can occur during library amplification. Again the snRNA-seq replicates show a stronger correlation between contaminating and endogenous counts, which can be explained by a stronger impact of the variation in capture efficiency and/or higher levels of barcode swapping.

**Figure 2.**
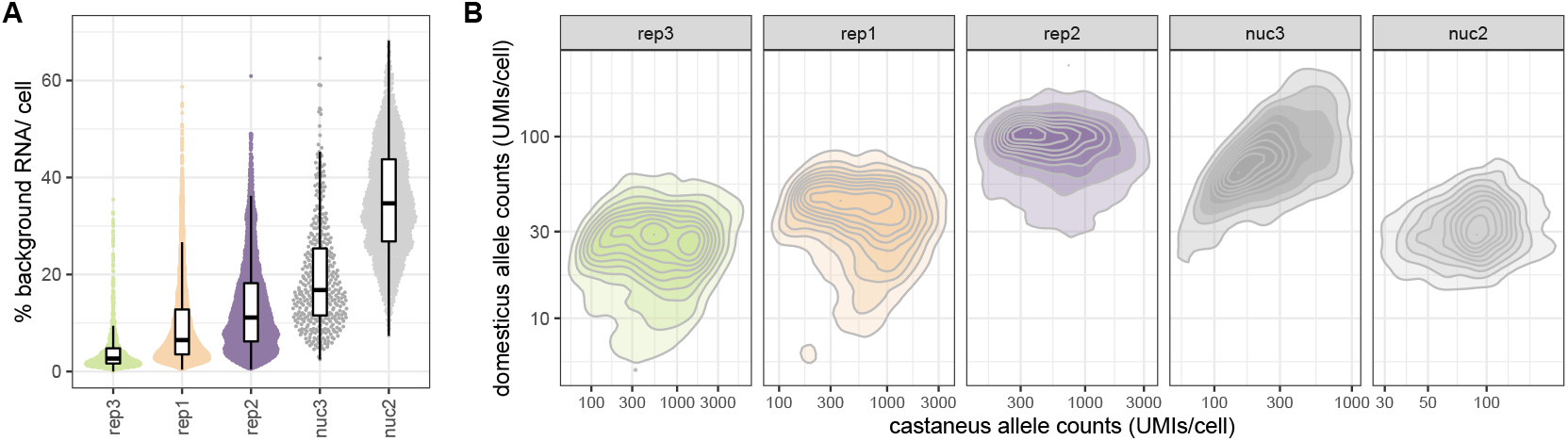
The level of background noise is variable across replicates and single cells. A) Estimated fraction of background noise per cell. The replicates on the x-axis are ordered by ascending median background noise fraction. B) In *M*.*m*.*castaneus* cells both endogenous *M*.*m*.*castaneus* specific alleles (x-axis) and *M*.*m*.*domesticus* specific alleles (y-axis) have coverage in each cell. The detection of *M*.*m*.*domesticus* specific alleles can be seen as background noise originating from cells of a different mouse.

However, by and large the absolute amount of background noise is approximately constant across cells and thus the contamination fraction mainly depends on the amount of endogenous RNA: the larger the cell, the smaller the fraction of background noise, pointing towards ambient RNA as the major source of the detected background (Figure 2B).

### The background noise profile does not always reflect the cell type composition

In order to better understand the effects of background noise, it is helpful to understand its origins and composition. To this end, we constructed pseudobulk profiles representing endogenous, contaminating and ambient expression profiles by using M. m. domesticus allele counts in *M. m. domesticus* cells (endogenous), *M. m. domesticus* allele counts in *M. m. castaneus* cells (contamination) and *M. m. domesticus* allele counts in empty droplets (empty) (Figure 3A, Supplementary Figure S4). In case of the three scRNA-seq replicates, we find that the contamination profiles correlate highly and similarly well with empty profiles (Spearman’s *ρ* = 0.73 − 0.85) and endogenous profiles (Spearman’s *ρ* = 0.70 − 0.87), while for the two snRNA-seq replicates the contamination profiles are clearly more similar to the empty (Spearman’s *ρ* ∼ 0.85) than to the endogenous profiles (Spearman’s *ρ* ∼ 0.50) (Figure 3B).

**Figure 3.**
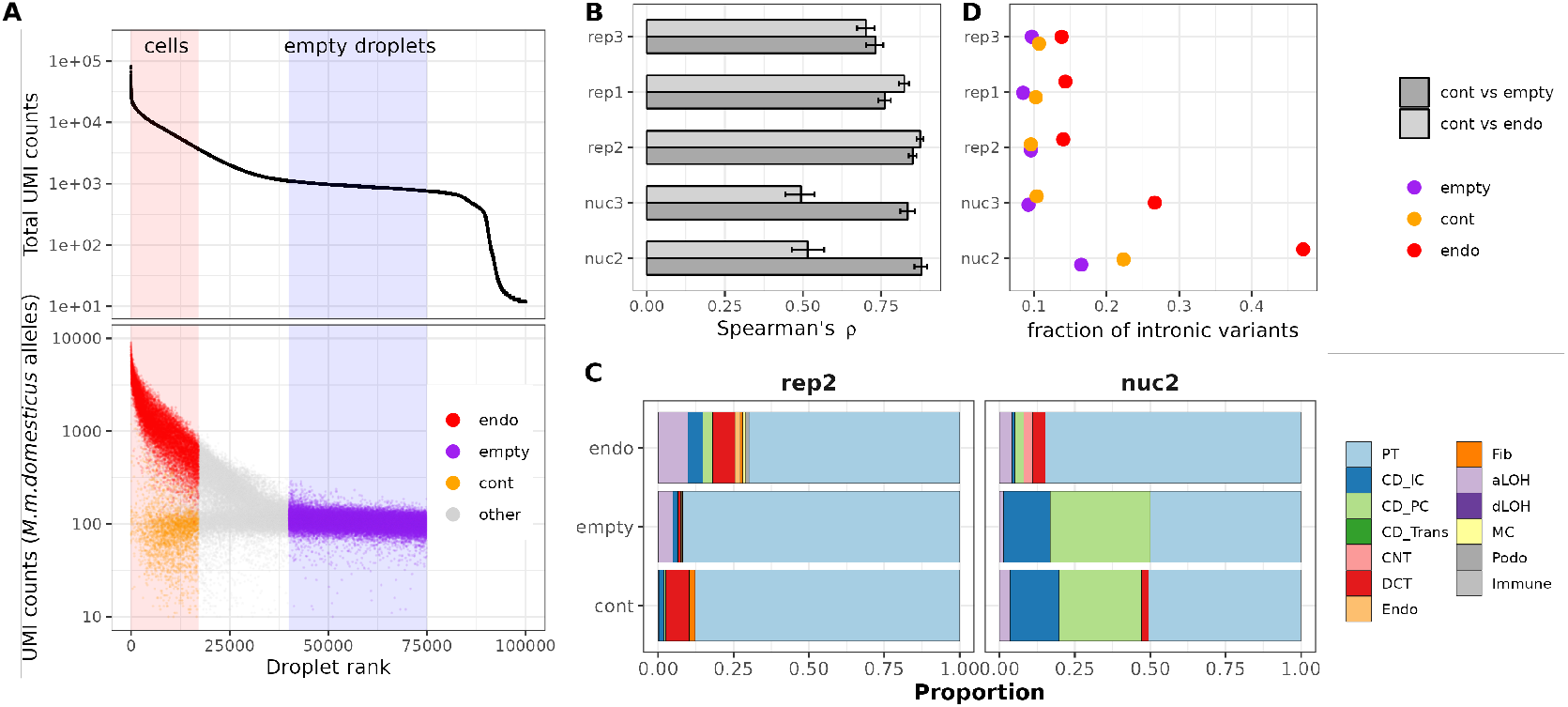
Characterization of ambient RNA in cells and empty droplets. A) Ordering droplet barcodes by their total UMI count to distinguish cell-containing droplets with high UMI counts from empty droplets that only contain cell-free ambient RNA and are identifiable as a plateau in the UMI curve, shown here for replicate 2. UMI counts of reads covering *M*.*m*.*domesticus* specific alleles were used to construct three profiles depending on whether they were associated with *M*.*m*.*domesticus* cell barcodes (endogenous counts, endo), *M*.*m*.*castaneus* cell barcodes (contaminating counts, cont) or empty droplet barcodes (empty). Counts from droplets that are not clearly assignable as cell-containing or empty were excluded from further analysis (other). B) Spearman rank correlation between pseudobulk profiles. C) Deconvolution of cell type contributions to each pseudobulk profile, exemplified by replicates rep2 and nuc2. The stacked barplots depict the estimated fraction of each cell type in the profile as inferred by SCDC using the annotated single cell data of each replicate as reference. PT: proximal tubule; CD_IC: intercalated cells of collecting duct; CD_PC: principal cells of collecting duct; CD_Trans: transitional cells of collecting duct; CNT: connecting tubule; DCT: distal convoluted tubule; Endo: endothelial; Fib: fibroblasts; aLOH: ascending loop of Henle; dLOH: descending loop of Henle; MC: mesangial cells; Podo: podocytes. D) Fraction of reads covering intronic variants in each of the three profiles.

Using deconvolution [16], we reconstructed the cell type composition of the pseudobulk profiles, and, in agreement with the correlation analysis, we find that in the scRNA-seq data the cell type compositions inferred for endogenous, contamination and empty counts are by and large similar with a slight increase in the PT-profile in empty droplets, suggesting that this cell type is more vulnerable to dissociation procedure than other cell types. In contrast, deconvolution of the empty droplet and contamination fraction of our snRNA-seq data, that in contrast to the scRNA-seq data were prepared from frozen samples, shows a clear shift in cell type composition with a decreased PT fraction (Figure 3C, Supplementary Figure S5).

Moreover, for the snRNA-seq data we expect that cytosolic mRNA contributes more to the contaminating profile than to the endogenous profile. Indeed, we find that in good nuclei (endogenous molecules) more than 25% of the allele counts fall within introns, while out of the molecules from empty droplets less than 18% fall within introns (Figure 3D). The intron fraction of the contaminating molecules lies in-between the endogenous and the empty droplet fraction, but is in all cases much closer to the empty intron fraction, thus suggesting again that the majority of the background noise likely originates from ambient RNA. However, the slight increase in the intron fraction of the contamination relative to empty droplets suggests that at least a small part of the observed background noise is due to barcode swapping.

### The impact of contamination on marker gene analyses

The ability to distinguish hitherto unknown cell types and states is one of the greatest achievements made possible by single cell transcriptome analyses. To this end, marker genes are commonly used to annotate cell clusters for which available classifications appear insufficient. An ideal marker gene would be expressed in all cells of one type but in none of the other present cell types. Thus, when comparing expression levels of one cell type versus all others, we expect high log2-fold changes, the higher the change the more reliable the marker. However, such a reliance on marker genes also makes this type of analysis vulnerable to background noise. Our whole kidney data can illustrate this problem well, because with the very frequent proximal tubular (PT) cells we have a dominant cell type for which rather specific marker genes are known [17]. Slc34a1 encodes a phosphate transporter that is known to be expressed exclusively in PT cells [18, 19]. As expected, it is expressed highly in PT cells, but it is also present in a high fraction of other cells (Figure 4A,E, Supplementary Figure S6). Moreover, the log2-fold changes of Slc34a1 are smaller in replicates with larger background noise, indicating that the detection of Slc34a1 in non-PT cells is likely due to contamination (Figure 4B-D). We observe the same pattern for other marker genes as well: they are detected across all cell types (Figure 4E, Supplementary Figure S7) and an increase of background noise levels goes along with decreasing log2-fold changes and increasing detection rates in other cell types (Figure 4F,G). Thus, the power to accurately detect marker genes decreases in the presence of background noise.

**Figure 4.**
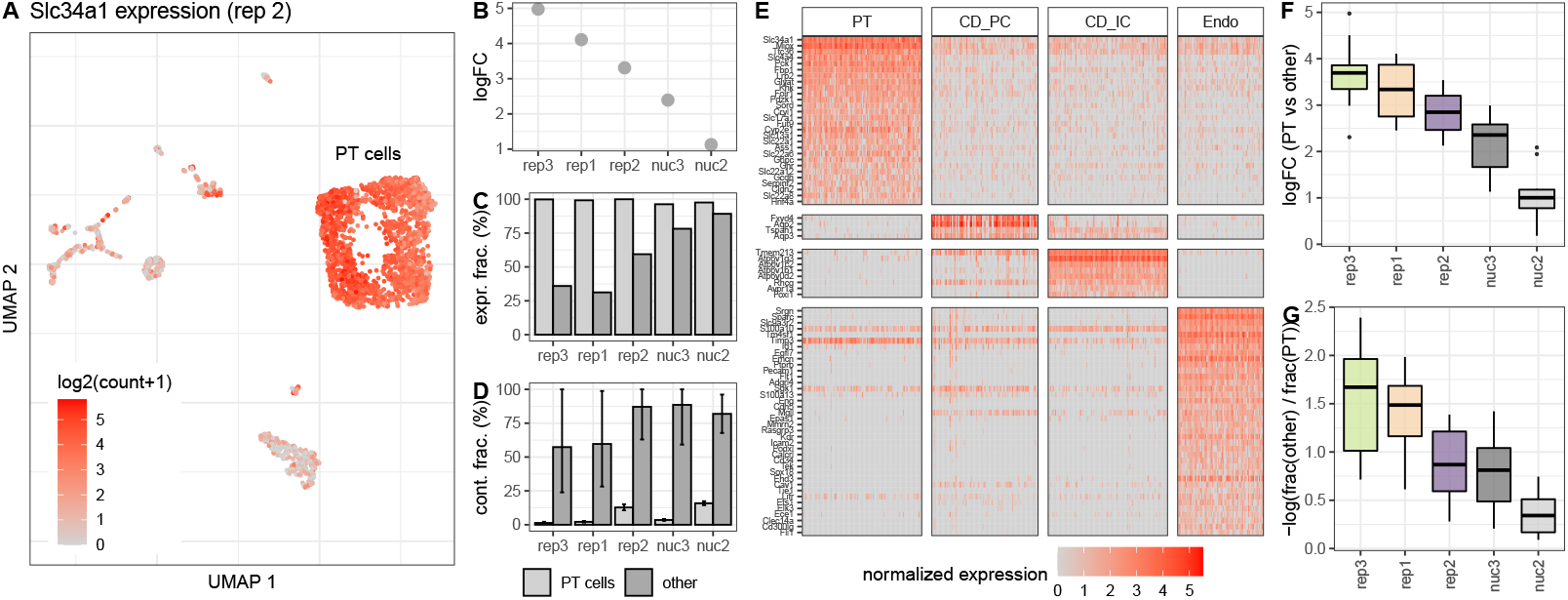
Background noise affects differential expression and specificity of cell type specific marker genes. A) UMAP representation of replicate 2 colored by the expression of Slc34a1, a marker gene for cells of the proximal tubule (PT). Besides high counts in a cluster of PT cells, Slc34a1 is also detected in other cell type clusters. Differential expression analysis between PT and all other cells shows a decrease of the detected log fold change of Slc34a1 (B) at higher background noise levels, as well as an increase of the fraction of non PT cells in which UMI counts of Slc34a1 were detected (C). D) Estimation of the background noise fraction of Slc34a1 expression indicates that the majority of counts in non PT cells originates from background noise. Error bars indicate 90% profile likelihood confidence intervals. E) Heatmap of marker gene expression for four cell types in replicate 2, downsampled to a maximum of 100 cells per cell type. F) Comparison across replicates of log2 fold changes of 10 PT marker genes calculated based on the mean expression in PT cells against mean expression in all other cells. G) For the same set of genes as in E), the log ratio of fraction of cells in which a gene was detected in others and PT cells shows how specific the gene is for PT cells.

### Benchmark of background noise estimation tools

Given that background noise will be present to varying degrees in almost all scRNA-seq and snRNA-seq replicates, the question is whether background removal methods can alleviate the problem without the information from genetic variants. SoupX [11], DecontX [16] and CellBender [4], all provide an estimate of the background noise level per cell. Here, we use our genotype-based background estimates as ground truth to compare it to the estimates of the three background removal methods (Figure 5A, Supplementary Figure S8). All methods have adjustable parameters, but also provide a set of defaults. For CellBender the user can adjust the nominal false positive rate to put a cap on losing information from true counts. For SoupX and DecontX the resolution of the clustering of cells that is later used to model the endogenous counts can be adjusted. In addition, SoupX can be provided with an expected background level and for DecontX the user can provide a custom background profile rather than using the default estimation strategy for the background profile. At least with our reference dataset, CellBender does not seem to profit from changing the defaults, while SoupX’s performance is boosted, if provided with realistic background levels (Supplementary Figure S13). Because in a real case scenario, the true background level is unknown, we decided to report the SoupX performance metrics under default settings. DecontX defaults to estimating the putative background profile from averaging across intact cells, but also gives the user the possibility to provide another profile, such as the profile of empty droplets as used in CellBender and SoupX. To ensure comparability, we report DecontX’s performance with empty droplets as background profile (DecontX_*background*_) in addition to DecontX with default settings (DecontX_*default*_).

**Figure 5.**
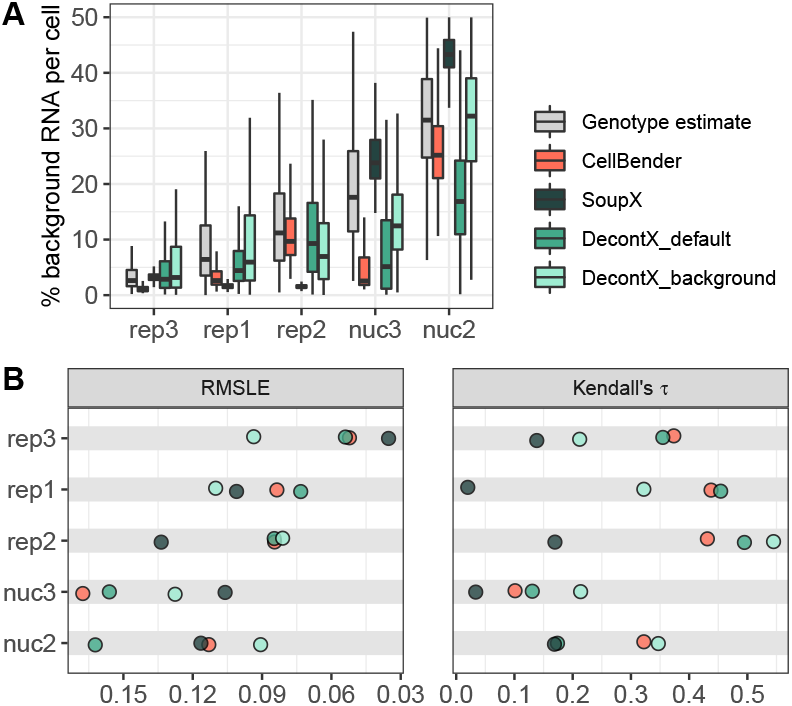
Accuracy of computational background noise estimation. A) Estimated background noise levels per cell based on genetic variants (grey) and different computational tools. B) Taking the genotype-based estimates as ground truth, Root Mean Squared Logarithmic Error (RMSLE) and Kendall rank correlation serve as evaluation metrics for cell-wise background noise estimates of different methods. Low RMSLE values indicate high similarity between estimated values and the assumed ground truth. High values of Kendall’s *τ* correspond to good representation of cell to cell variability in the estimated values.

We find that CellBender and DecontX can estimate background noise levels similarly well for the scRNA-seq replicates, while SoupX tends to underestimate background levels and also cannot capture the cell to cell variation as measured by the correlation with the ground truth (Figure 5B). For the snRNA-seq data, SoupX performs better at estimating global background levels, but as for the scRNA-seq still cannot capture cell to cell variation. In contrast, both CellBender and DecontX perform worse with the snRNA-seq data than with the scRNA-seq data. In the case of DecontX, the default setting provides much worse estimates than the estimates using empty droplets as background profile.

All in all, CellBender shows the most robust performance across replicates with default settings, while DecontX’ and SoupX’ performance seems to require parameter tuning. In the case of DecontX the default works well for scRNA-seq data, but not for snRNA-seq data, while for SoupX the opposite is true.

A drawback of CellBender is its runtime. While SoupX and DecontX take seconds and minutes to process one 10x channel, CellBender takes ∼ 45 CPU hours. However, parallelization is possible.

All methods struggle most with the nuc3 replicate that has the fewest *M*.*m. castaneus* cells and the lowest cell type diversity among our five data sets (Figure 1B,E). This also presents a problem for other downstream analyses and thus we do not consider nuc3 further.

### Effect of background noise removal on marker gene detection

Above we have shown that computational methods can estimate background noise levels per cell. Moreover, all three methods provide the user with a background corrected count matrix for downstream analysis. Here, we compare the outcomes of marker gene detection, clustering and classification when using corrected count matrices from SoupX, DecontX and CellBender (Figure 6A, Supplementary Figure S9). To characterize the impact on marker gene detection, we first check in how many cells an unexpected marker gene was detected; for example, how often Slc34a1 was detected in cells other than PTs (Figure 6B). Without correction we find Slc34a1 reads in ∼ 60% of non-PT cells of scRNA-seq rep2, SoupX reduces this rate to 54%, CellBender to 7% and DecontX_*background*_ to 9%. DecontX_*default*_ manages to remove most contaminating reads reducing the Slc34a1 detection rate outside PTs to 2%. While we find a similar ranking when averaging across several marker genes from the PanglaoDB database [17] and scRNA-seq replicates (Figure 6C), the ranking changes for nuc2: DecontX_*default*_ fails: after correction, Slc34a1 is still found in 87% of non-PT cells while DecontX_*background*_ is better with a rate of 20%. Here, CellBender and SoupX are clearly better with reducing the Slc34a1 detection rate to 4% and *<* 1%, respectively (Supplementary Figure S10).

**Figure 6.**
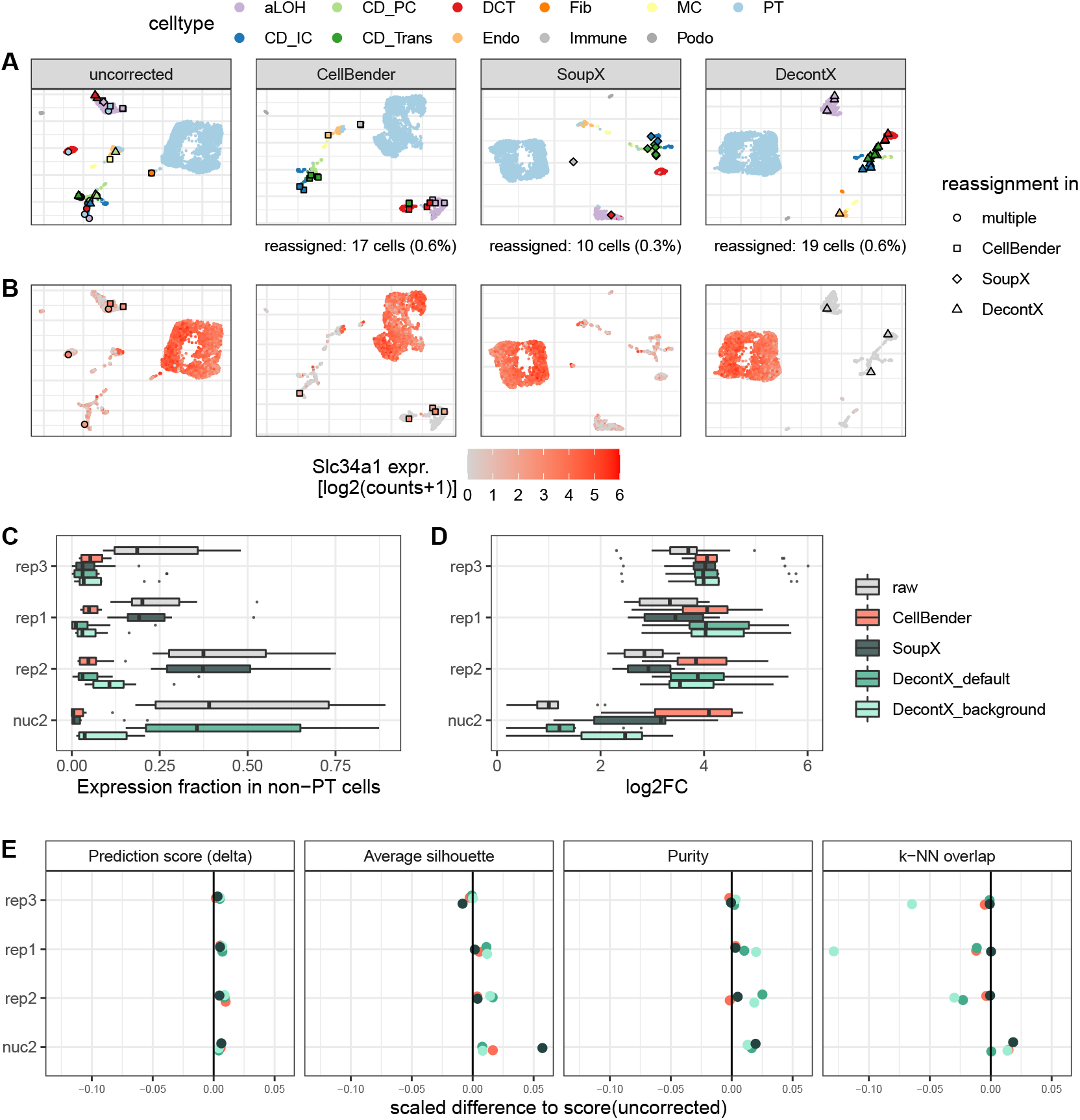
Effect of Background removal on downstream analysis. A) UMAP representation of replicate 2 single-cell data before and after background noise correction, colored by cell type labels obtained from reference based classification. Individual cells that received a new label after correction are highlighted. PT: proximal tubule; CD_IC: intercalated cells of collecting duct; CD_PC: principal cells of collecting duct; CD_Trans: transitional cells of collecting duct; CNT: connecting tubule; DCT: distal convoluted tubule; Endo: endothelial; Fib: fibroblasts; aLOH: ascending loop of Henle; dLOH: descending loop of Henle; MC: mesangial cells; Podo: podocytes B) Expression of the PT cell marker Slc34a1 before and after background noise correction in replicate 2. Cells that were classified as PT cells in the uncorrected data, but got reassigned after correction, are highlighted. C),D) Differential expression analysis of 10 PT markers, evaluating the expression fraction in non-PT cells (C) and the log2 fold change between PT and all other cells (D). E) Evaluation metrics for the effect of background noise correction on classification and clustering. For each metric the change relative to the uncorrected data is depicted. The values were scaled by the possible range of each metric. Prediction score: cell-wise score “delta” of reference based classification with SingleR [21]. Average silhouette: Mean of silhouette widths per cell type. Purity: Cluster purity calculated on cell type labels as ground truth and Louvain clusters as test labels. *k* -NN overlap: overlap of the *k* =50 nearest neighbors per cell compared to genotype-cleaned reference *k* -NN graph.

Even though the changes in the marker gene detection rates outside the designated cell type seem dramatic, with moderate background levels as e.g. in rep2, the identification of marker genes [20] is affected only a little. CellBender correction has the largest effect on marker gene detection, yet 8 from the top 10 genes without correction remain marker genes with CellBender correction (Spearman’s correlation for top 100 *ρ* = 0.84). In contrast, in the nuc2 data with high background levels, the change in marker gene detection is dramatic. Here, only one of the top 10 marker genes remains after correction (Spearman’s correlation for top 100 *ρ* = 0.04). The largest improvement is achieved with CellBender: After correction, four out of the top 10 were known marker genes [17], while this overlap amounted to only one in the raw data (Supplementary Figure S11B). Moreover, we find that background removal also increases the detected log-fold-changes of known marker genes across all replicates and methods, with CellBender providing the largest improvement (Figure 6D, Supplementary Figure S11C).

### Effect of background noise removal on classification and clustering

One of the first and most important tasks in single cell analysis is the classification of cell types. As described above, we could identify 13 cell types in our uncorrected data using an external single cell reference dataset [14, 21]. Going through the same classification procedure after correction for background noise, changes the classification of only very few cells (Figure 6A, Supplementary Figure S9). For the scRNA-seq experiments *<* 1% and for the snRNA-seq data up to 1.3% of cells change labels after background removal compared to the classification using raw data. Before correction, these cells are mostly located in clusters dominated by a different cell type (Figure 6A). Moreover, these cells tend to have higher background levels as exemplified by the PT-marker gene Slc34a1 (Figure 6B). Finally, background removal - irrespective of the method - improves the classification prediction scores (Figure 6E, Supplementary Figure S12). Together, this indicates that background removal improves cell classification.

Similarly, background removal also results in more distinct clusters. Here, we reason that cells of the same cell type should cluster together and evaluate the impact of background removal 1) on the silhouette scores for cell types and 2) on the cell type purity of each cluster using unsupervised clustering (Figure 6E). For the scRNA-seq data DecontX results in the purest and most distinct clusters, while for the snRNA-seq data SoupX wins in these categories.

All in all, it seems clear that all background removal methods sharpen the broad structure of the data a little, but how about fine structure? To answer this question, we turn again to the genotype cleaned data to obtain a ground truth for the *k* -nearest neighbors of a cell and calculate how much higher the overlap of the background corrected data is with this ground truth as compared to using the raw data (Figure 6E). For the scRNA-seq data, DecontX has the largest improvement on the broad structure, but at same time in particular DecontX_*background*_ lowers the overlap in *k* -NN with our assumed ground truth, suggesting that this change in structure is a distortion rather than an improvement. SoupX leaves the fine structure by and large unchanged in the scRNA-seq data, while both CellBender and DecontX make the fine structure slightly worse. In contrast, for the high background levels of the nuc2, all background removal methods achieve an improvement, with SoupX and CellBender performing best.

## Discussion

Here we provide a dataset for the characterization of background noise in 10x Genomics data that is ideal to benchmark background removal methods. The mixture of cell types in our kidney data provides us with realistic cell type diversity and the mixture of mouse subspecies enables us to identify foreign alleles in a cell, thus resulting in a dataset that allows us to quantify background noise across diverse cell types and features. Moreover, the replicates have very different contamination levels, making it possible to assess the impact of low, intermediate and high background levels. As expected, marker gene identification is affected and markers appear less specific, as they are detected in cell types where they are not expressed. The severity of the issue directly depends on background noise levels (Figure 4). This particular problem has been observed previously and has been used as a premise to develop background correction methods [22, 4, 11].

The novelty of this analysis is that - thanks to the mix of mouse subspecies - we are able to obtain expression profiles that describe the source of contamination in each sample and also have a ground truth for a more realistic dataset. We started to characterize background noise by comparing the contamination profile with the profile of empty droplets and that of endogenous counts of good cells. In agreement with the idea that ambient RNA is due to leakage of cytosol, we find that empty droplets show less evidence for unspliced mRNA molecules and that the unspliced fraction in the contamination profiles is similar to that of empty droplets, suggesting that the majority of the detected background noise is due to ambient RNA. Only in the snRNA-seq dataset nuc2 the unspliced fraction of the contamination profile is clearly higher than for empty droplets, providing evidence for at least some barcode swapping (Figure 3C). Hence, the observed correlation between cell size and the absolute amounts of background noise per cell in most of the replicates is likely due to variation in dropout rates [4] (Figure 2B).

Another important insight from comparing contamination, empty and endogenous profiles is that we can deduce the origin of the contamination. While for the scRNA-seq data all three profiles are highly correlated and are the result of very similar cell type mixtures, for the snRNA-seq data the empty and the contamination profiles are distinct from the expected endogenous mixture profile. Encouragingly the endogenous profile of the snRNA-seq data agrees well with the cell type proportions from our scRNA-seq data as well as the literature [14, 23], suggesting that neither library preparation method introduces a strong cell type bias. Moreover, the higher similarity between the empty and the contamination profiles strongly supports again that the majority of background noise is ambient RNA and hence using the empty rather than the endogenous profile as a reference to model background noise is a good choice. Indeed, the performance of DecontX for nuc2 is improved by providing the empty droplet profile as compared to the endogenous profile which is the default (Figure 5A). We also observed that SoupX performs much better for the snRNA-seq data than the scRNA-seq data. We speculate that the marker gene identification that is the basis for estimating the experiment-wide average contamination is hampered by the fact that our dataset has one very dominant cell type that has the same prevalence in the empty droplets, thus masking all background. However, even if SoupX gets the overall background levels right, it by design grossly underestimates the variance among cells and cannot capture the cell to cell variation (Figure 5B,C). Overall CellBender provides the most accurate estimates of the background noise levels and also captures the cell to cell variation rather well.

In line with this, marker gene detection is most improved by CellBender, which is the only method that removes marker gene molecules from other cell types and increases the log-fold-change consistently well. The effect of background removal on other downstream analyses is much more subtle. For starters, classification using an external reference is rather robust. Even with high levels of background noise, background removal improves classification only for a handful of cells and we cannot say that one method outperforms the others (Figure 6E, Supplementary Figure S12). Similarly, the broad structure of the data improves only minimally and this minimal improvement comes at the cost of disrupting fine structure (Figure 6E). Here, again CellBender strikes the best balance between removing variation but preserving the fine structure, while DecontX tends to remove too much within-cluster variability, as the *k* -NN overlap with the genotype-based ground truth for DecontX is even lower than for the raw data. All in all, CellBender shows the best performance in removing background noise.

## Conclusion

Levels of background noise can be highly variable within and between replicates and the contamination profiles do not always reflect the cell type proportions of the sample. Marker gene detection is affected most by this issue, in that known cell type specific marker genes can be detected in cell clusters where they do not belong. Existing methods for background removal are good at removing such stray marker gene molecule counts. In contrast, classification and clustering of cells is rather robust even at high levels of background noise. Consequently, background removal improves the classification of only few cells. Moreover, it seems that for low and moderate background levels the tightening of existing broad structures may go at the cost of fine structure. In summary, for marker gene analysis, we would always recommend background removal, but for classification, clustering and pseudotime analyses, we would only recommend background removal when background noise levels are high.

## Methods

### Mice

Three mouse strains were ordered from Jackson Laboratory at 6-8 weeks of age: C57BL/6J (000664), CAST/EiJ (000928), and 129S1/SvlmJ (002448). All animals were subjected to intracardiac perfusion of PBS to remove blood. Kidneys were dissected, divided into 1/4s, and subjected to the tissue dissociation protocol, stored in RNAlater, or snap-frozen in liquid nitrogen.

### Tissue dissociation for single cell isolation

The single cell suspensions were prepared following an established protocol [24] with minor modifications. In detail, one of each kidney sagittal quarter from three perfused mice of different strains C57BL/6, CAST/EiJ and 129S1/SvImJ were harvested into cold RPMI (Thermo Fisher Scientific, 11875093) with 2% heat-inactivated Fetal Bovine Serum (Gibco, Thermo Fisher Scientific, 16140-071; FBS) and 1% penicillin/streptomycin (Gibco, Thermo Fisher Scientific, 15140122). Each piece of the tissue was then minced for 2 minutes with a razor blade in 0.5 ml 1x liberase TH dissociation medium (10x concentrated solution from Millipore Sigma, 05401135001, reconstituted in DMEM/F12(Gibco, Thermo Fisher Scientific, 11320-033 in a petri dish on ice. The chopped tissue pieces were then pooled into one 1.5 ml Eppendorf tube and incubated in a thermomixer at 37°C for 1 hour at 600rpm with gentle pipetting for trituration every 10 minutes. The digestion mix was then transferred to a 15 ml conical tube and mixed with 10 ml 10% FBS RPMI. After centrifugation in a swinging bucket rotor at 500g for 5 min at 4°C and supernatant removal, the pellet was resuspended in 1ml red blood cell lysing buffer (Sigma Aldrich, R7757). The suspension was spun down at 500g for 5 min at 4°C followed by supernatant removal. The pellet cleared of the red blood cell ring was then resuspended in 250 μl Accumax (Stemcell Technologies, 7921) and incubated at 37°C for 3 mins. The reaction was stopped by mixing with 5 ml 10% FBS RPMI and spinning down at 500g for 5 min at 4°C followed by supernatant removal. The cell pellet was then resuspended in PBS with 0.4% BSA (Sigma, B8667) and passed through a 30 μm filter (Sysmex, 04-004-2326). The cell suspension was then assessed for viability and concentration using the K2 Cellometer (Nexcelom Bioscience) with the AOPIcell stain (Nexcelom Bioscience, CS2-0106-5ML).

### Nuclei isolation from RNAlater preserved frozen tissue

The single nuclei suspensions were prepared following an established protocol [25] with minor modifications. In detail, the RNALater reserved frozen tissue of 3 mice kidney quarters were thawed and transferred to one petri dish preloaded with 1 ml TST buffer containing 10 mM Tris, 146 mM NaCl, 1 mM CaCl2, 21 mM MgCl2, 0.03% Tween-20 (Roche, 11332465001) and 0.01% BSA (Sigma, B8667). It was minced with a razor blade for 10 minutes on ice. The homogenized tissue was then passed through a 40 μm cell strainer (VWR, 21008-949) into a 50 ml conical tube. One ml TST buffer was used to rinse the petri dish and collect the remaining tissue into the same tube. It was then mixed with 3 ml of ST buffer containing 10 mM Tris, 146 mM NaCl, 1 mM CaCl2 and 21 mM MgCl2 and spun down at 500g for 5 min at 4°C followed by supernatant removal. In the second experiment this washing step was repeated 2 more times. The pellet was resuspended in 100 μl ST buffer and passed through a 35 μm filter. The nuclei concentration was measured using the K2 Cellometer (Nexcelom Bioscience) with the AO nuclei stain (Nexcelom Bioscience, CS1-0108-5ML).

### Single-cell and single-nucleus RNA-seq

The cells or nuclei were loaded onto a 10x Chromium Next GEM G chip (10x Genomics, 1000120) aiming for recovery of 10,000 cells or nuclei. The RNA-seq libraries were prepared using the Chromium Next GEM Single Cell 3’ Reagent kit v3.1 (10x Genomics, 1000121) following vendor protocols. The libraries were pooled and sequenced on NovaSeq S1 100c flow cells (Illumina) with 28 bases for read1, 55 bases for read2 and 8 bases for index1 and aiming for 20,000 reads per cell.

### Processing and annotation of scRNA-seq and snRNA-seq data

The scRNA-seq and snRNA-seq data were processed using Cell Ranger 3.0.2 using as reference genome and annotation mm10 version 2020A for the scRNA-seq data and and a pre-mRNA version of mm10 2.1.0 as reference for snRNA-seq. In order to identify cell containing droplets we processed the raw UMI matrices with the DropletUtils package [5]. The function barcodeRanks was used to identify the inflection point on the total UMI curve and the union of barcodes with a total UMI count above the inflection point and Cell Ranger cell call were defined as cells.

For cell type assignment we used 3 scRNA-seq and 4 snRNA-seq experiments from Denisenko et al. [14] as a reference. Cells labeled as “Unknown” (n=46), “Neut” (n=17) and “Tub” (n=1) were removed. The reference was log-normalized and split into seven count matrices based on chemistry, preservation and dissociation protocol. Subsequently, a multi-reference classifier was trained using the function *trainSingleR* with default parameters of the R package SingleR version 1.8.1 [21]. After this processing, we could use the data to classify our log-normalized data using the *classifySingleR* function without fine-tuning (fine.tune = F). Hereby, each cell is compared to all seven references and the label from the highest-scoring reference is assigned. Some cell type labels were merged into broader categories after classification: cells annotated as “CD_IC”, “CD_IC_A” or “CD_IC_B” were relabeled as “CD_IC”, cells annotated as “T”, “NK”, “B” or “MPH” were relabeled as “Immune”. Cells that were unassigned after pruning of assignments based on classification scores were removed for subsequent analyses.

### Demultiplexing of mouse strains

A list of genetic variants between mouse strains was downloaded in VCF format from the Mouse Genomes Project [13], accessed on 21 October 2020. This reference VCF file was filtered for samples CAST_EiJ, C57BL_6NJ and 129S1_SvImJ and chromosomes 1-19. Genotyping of single barcodes was performed with cellsnp-lite [26], filtering for positions in the reference VCF with a coverage of at least 20 UMIs and a minor allele frequency of at least 0.1 in the data (–minCOUNT 20, –minMAF 0.1). Vireo [22] was used to demultiplex and label cells based on their genotypes. Only cells that could unambigously assigned to CAST_EiJ (CAST), C57BL_6NJ (BL6) or 129S1 SvImJ (SvImJ) were kept, cells labeled as doublet or unassigned were removed.

### Genotype-based estimation of background noise

Based on the coverage filtered VCF-file (see above), we identified homozygous SNPs that distinguish the three strains and removed SNPs that had predominantly coverage in only one of the strains (1st percentile of allele frequency).

In most parts of the analysis, we focused on the comparison between the mouse subspecies, *M*.*m*.*domesticus* and *M*.*m*.*castaneus*. To this end, we subseted reads (UMI-counts) that overlap with SNPs that distinguish the two mouse subspecies.

To estimate background noise levels based on allele counts of genetic variants, an approach described in Heaton et al.[15] was adapted to estimate the total amount of background noise for each cells. First, the abundance of endogenous and foreign allele counts (i.e. cross-genotype background noise) was quantified per cell. Because of the filter for homozygous variants, there are two possible genotypes for each locus, denoted as 0 for the endogenous allele, i.e. the expected allele based on the strain assignment of the cell, and 1 for the foreign allele. The probability for observable background noise at each locus *l* in cell *c* is given by

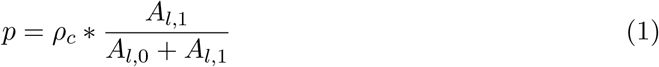

where *ρ*_*c*_ is the total background noise fraction in a cell and the experiment wide (over cells and empty droplets) foreign allele fraction is calculated from the foreign allele counts *A*_*l*,1_ and the endogenous allele counts *A*_*l*,0_. The foreign allele fraction is then used to account for intra-genotype background noise (contamination within endogenous allele counts).

The observed allele counts *A*_*c*_ per cell are modeled as draws from a binomial distribution with the likelihood function:

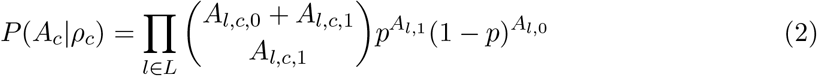

A maximum likelihood estimate of *ρ*_*c*_ was obtained using one dimensional optimization in the interval [0,1].

The 95% confidence interval of each *ρ*_*c*_ estimate was calculated as the profile likelihood using the function *uniroot* of the R package stats [27].

### Comparison of endogenous, contamination and empty droplet profiles

Empty droplets were defined based on the UMI curve of the barcodes ranked by UMI counts, thus selecting barcodes from a plateau with ∼ 500 − 1000 UMIs (Supplementary Figure S4). For the following analysis, the presence of *M*.*m*.*domesticus* alleles in *M*.*m*.*domesticus* cells (i.e., endogenous), in *M*.*m*.*castaneus* cells (i.e., contamination) and empty droplets was compared. After this filtering, we summarized counts per gene and across barcodes of the same category to generate pseudobulk profiles.

In order to estimate cell type composition in the empty and contamination profiles, we used the deconvolution method implemented in SCDC[16], the endogenous single cell allele counts from the respective replicate were used as reference (*qcthreshold=0*.*6*). In addition, cell type filtering (frequency*>*0.75%) was applied. Endogenous, contamination and empty pseudobulk profiles from each replicate were deconvoluted using their respective single cell / single nucleus reference.

To compare the correlation between the different profiles, pseudobulk counts were downsampled to the same total size.

### Evaluation of marker gene expression

A list of marker genes for Proximal tubule cells (PT), Principal cells (CD PC), Intercalated cells (CD_IC) and Endothelial cells (Endo) was downloaded from the public database PanglaoDB [17], accessed on 13 May 2022. Log2 fold changes contrasting PT cells against all other cells were calculated with Seurat using the function *FindMarkers* after normalization with *NormalizeData*. The expression fraction *e* of PT markers was calculated as the fraction of cells for which at least 1 count of that gene was detected. To contrast expression fraction in PT cells against non-PT, the negative log-ratio was calculated as −*log*((*e*_*P T*_ + 1)*/*(*e*_*non*−*P T*_ + 1)).

### Computational background noise estimation and correction methods

**CellBender** [4] makes use of a deep generative model to include various potential sources of background noise. Cell states are encoded in a lower-dimensional space and an integer matrix of noise counts is inferred, which is subsequently subtracted from the input count matrix to generate a corrected matrix.

The *remove-background* module of CellBender v0.2.0 was run on the raw feature barcode matrix as input, with a default *fpr* value of 0.01. For the comparison of different parameter settings, *fpr* values of 0.05 and 0.1 were also included in the analysis. For the parameter *expected-cells* the number of cells after cell calling and filtering in each replicate was provided. The parameter *total-droplets-included* was set to 25000.

**SoupX**[11] estimates the experiment-wide amount of background noise based on the expression of strong marker genes that are expected to be expressed exclusively in one cell type. These genes can either be provided by the user or identified from the data. A profile of background noise is inferred from empty droplets. This profile is subsequently removed from each cell after aggregation into clusters to generate a corrected count matrix.

Cluster labels for SoupX were generated by Louvain clustering on 30 principal components and a resolution of 1 as implemented by *FindClusters* in Seurat after normalization and feature selection of 5000 genes. Providing the CellRanger output and cluster labels as input, data were imported into SoupX version 1.6.1 and the background noise profile was inferred with *load10X*. The contamination fraction was estimated using *autoEstCont* and background noise was removed using *adjustCounts* with default parameters.

For the comparison of parameter settings, different resolution values (0.5,1,2) for Lou-vain clustering were tested, alongside with manually specifying the contamination fraction (0.1,0.2).

**DecontX**[8] is a Bayesian method that estimates and removes background noise by modeling the expression in each cell as a mixture of multinomial distributions, one native distribution cell’s population and one contamination distribution from all other cell populations. The main inputs are a filtered count matrix only containing barcodes that were called as cells and a vector of cluster labels. The contamination distribution is inferred as a weighted combination of multiple cell populations. Alternatively, it is also possible to obtain an empirical estimation of the contamination distribution from empty droplets in cases where the background noise is expected to differ from the profile of filtered cells. The function *decontX* from the R package celda version 1.12.0 was run on the filtered, unnormalized count matrix and clusters were inferred with the implemented default method based on UMAP dimensionality reduction and dbscan [28] clustering. For the “DecontX_default” results the parameter’background’ was set to NULL, i.e. estimating background noise based on cell populations in the filtered data only. “DecontX_background” results were obtained by providing an unfiltered count matrix including all detected barcodes as’background’ to empirically estimate the contamination distribution. Besides the default clustering method implemented in DecontX, cluster labels obtained from Lovain clustering (resolution 0.5,1 and 2) were also provided to test different parameter settings.

### Evaluation metrics

#### Estimation accuracy

The genotype-based estimates *ρ*_*c*_ for *M*.*m*.*castaneus* cells served as ground truth to evaluate the estimation accuracy of different methods. For each method cell-wise background noise fractions *a*_*c*_ were calculated from the corrected count matrix *X* and the uncorrected (”raw”) count matrix

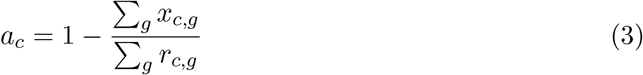

for cells *c* and genes *g*.

#### RMSLE

The Root Mean Squared Logarithmic Error (RMSLE) is a lower bound metric that we use to quantify the difference between estimated background noise fractions per cell *a*_*c*_ from different computational background correction methods and the genotype-based estimates *ρ*_*c*_, obtained from genotype based estimation. It is calculated as:

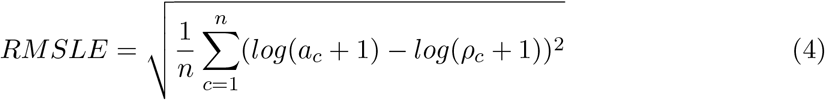

#### Kendall’s *τ*

To evaluate how well cell-to-cell variation of the background noise fraction is captured by the estimated values *a*_*c*_, the Kendall rank correlation coefficient *τ* to the genotype-based estimates *ρ*_*c*_ was computed using the implementation in the R package stats [27] as *τ* = *cor*(*a*_*c*_, *ρ*_*c*_, *method* = “*kendall*”).

#### Marker gene detection

The same set of 10 PT marker genes from PanglaoDB as in section Evaluation of marker gene expression was used to evaluate the improvement on marker gene detection on corrected count matrices.

**Log2 fold change** for each gene between the average expression in PT cells and average expression in other cells were obtained using the *NormalizeData* and *FindMarkers* functions in Seurat version 4.1.1.

#### Expression fraction

Entries in each corrected count matrix were first rounded to the nearest integer. The expression fraction of each gene in a cell population was calculated as the fraction of cells for which at least 1 count of that gene was detected. For evaluation of PT marker genes, unspecific detection is defined as the expression fraction in non-PT cells.

### Cell type identification

#### Prediction score

Each corrected count matrix was log-normalized and reference-based classification in SingleR [21] was performed with a pre-trained model (see section Processing and annotation of scRNA-seq and snRNA-seq data) on data from Denisenko et al. [14]. SingleR provides *delta* values as a measure for classification confidence, which depicts the difference of the assignment score for the assigned label and the median score across all labels. The *delta* values for each cell were retrieved using the function *getDeltaFromMedian* relative to the cells highest-scoring reference. A prediction score per cell type was calculated by averaging *delta* values across individual cells and a global prediction score per replicate was calculated by averaging across cell type prediction scores.

#### Average silhouette

The silhouette width is an internal cluster evaluation metric to contrast similarity within a cluster with similarity to the nearest cluster. The cell type annotations from reference-based classification were used as cluster labels here. Count matrices were filtered to select for *M*.*m*.*castaneus* cells and cell types with more than 10 cells. Distance matrices were computed on the first 30 principal components using euclidean distance as distance measure. Using the cell type labels and distance matrix as input, the average silhouette width per cell type was computed with the R package cluster version 2.1.4. An *Average silhouette* per replicate was calculated as the mean of cell type silhouette widths.

**Purity** is an external cluster evaluation metric to evaluate how well a clustering recovers known classes. Here, *Purity* was used to assess to what extent unsupervised cluster labels correspond to cell types. Count matrices were filtered to select for *M*.*m*.*castaneus* cells and cell types with more than 10 cells and louvain clustering as implemented in *FindClusters* of Seurat version 4.1.1 on the first 30 principal components and with a resolution parameter of 1 was used get a cluster label for each cell. Providing cell type annotations as true labels alongside the cluster labels, *Purity* was computed with the R package ClusterR version 1.2.6 [29].

#### *k* -NN overlap

To evaluate the lower-dimensional structure in the data beyond clusters and cell-types *k* -NN overlap was used as described in Ahlmann-Eltze and Huber [30]. A ground truth reference *k* -NN graph was constructed on a’genotype-cleaned’ count matrix, only counting molecules that carry a subspecies-endogenous allele. Raw and corrected count matrices were filtered to contain the same genes as in the reference and a query *k* -NN graph was computed on the first 30 principal components. The *k* -NN overlap summarizes the overlap of the 50 nearest neighbors of each cell in the query with the reference *k* -NN graph.

## Abbreviations

CAST: Mus musculus castaneus
*k*-NN: *k* nearest neighbor
snRNA-seq: single nucleus RNA-sequencing
PT: proximal tubular cells/markers
scRNA-seq: single cell RNA-sequencing
SNP: single nucleotide polymorphism
UMI: unique molecular identifier
UMAP-: Uniform Manifold Approximation and Projection
VCF: Variant Call Format

## 1 Declarations

### Ethics approval and consent to participate

All procedures performed are IACUC approved on Broad Institute animal protocol # 0061-07-15-1.

### Consent for publication

Not applicable.

### Availability of data and materials

The code used to analyse the data and benchmark the background methods is available on github https://github.com/Hellmann-Lab/scRNA-seq Contamination, larger files are deposited in the linked zenodo account. All sequencing files were deposited in GEO SRPXXXX.

### Author’s contributions

IH, WE and PJ conceptualized this study. IH and PJ wrote the original draft. PJ, BV and ZK conducted the formal analysis. SS did the data curation. XA, JM and CM performed the experiments. JL supervised the experiments. WE, HH, and JL acquired funding. All authors reviewed and edited the manuscript. (using https://credit.niso.org/)

## Acknowledgements

We thank Gabriela Stumberger for her help in benchmarking and Batuhan Akçabozan for his contribution to calculating genotype estimates. We thank the Broad Genomics Platform for sequencing.

## Supplementary Information

**Supplementary Figure S1.**
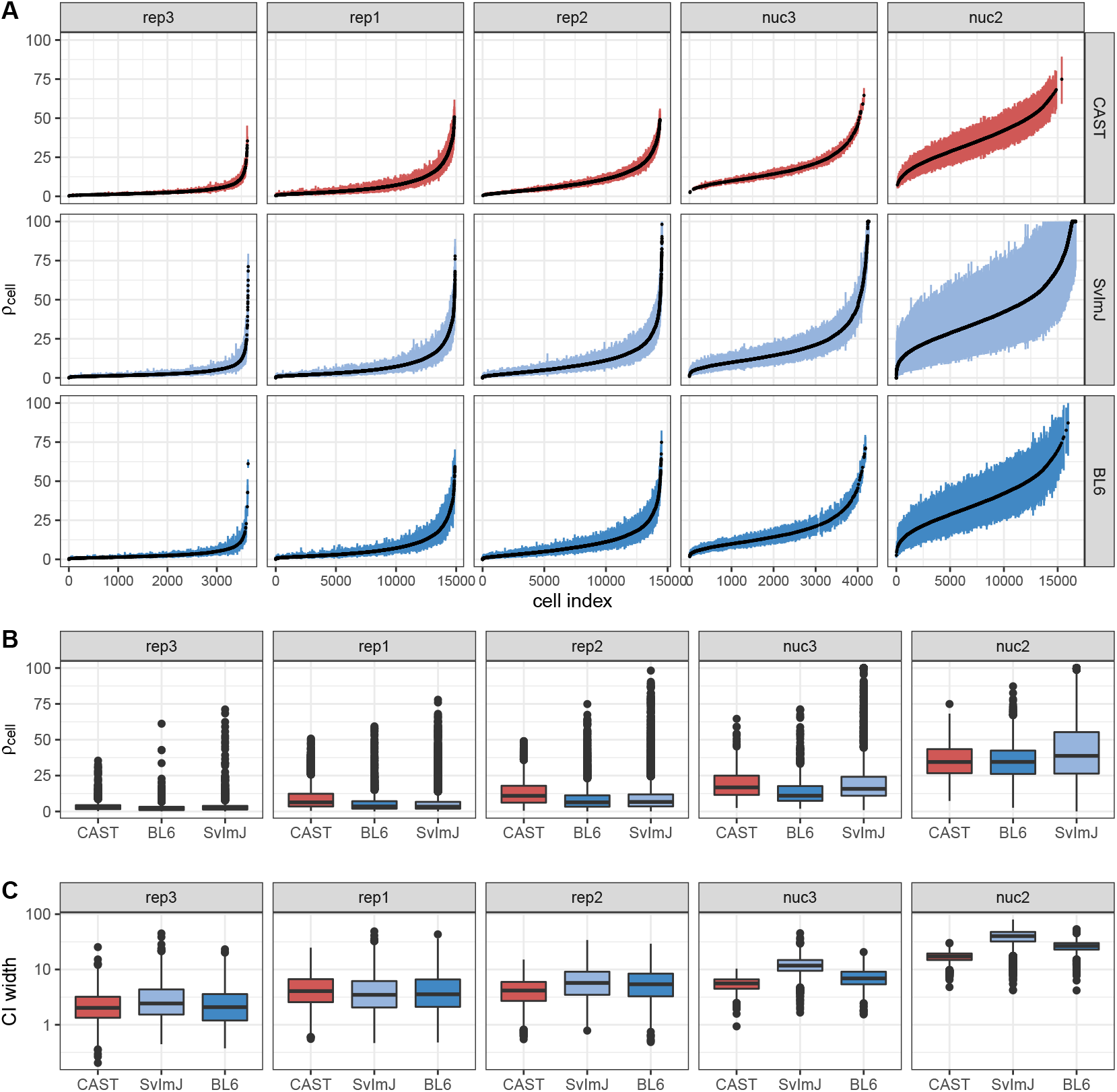
Estimation of background noise levels. A) Estimates of background noise (*ρ*_*cell*_) per cell. Cells were ordered by ascending *ρ*_*cell*_ in each replicate. Colored bars indicate 95% confidence intervals calculated by profile likelihood. B) Summary of *ρ*_*cell*_ estimates per strain. C) Width of confidence intervals for *ρ*_*cell*_.

**Supplementary Figure S2.**
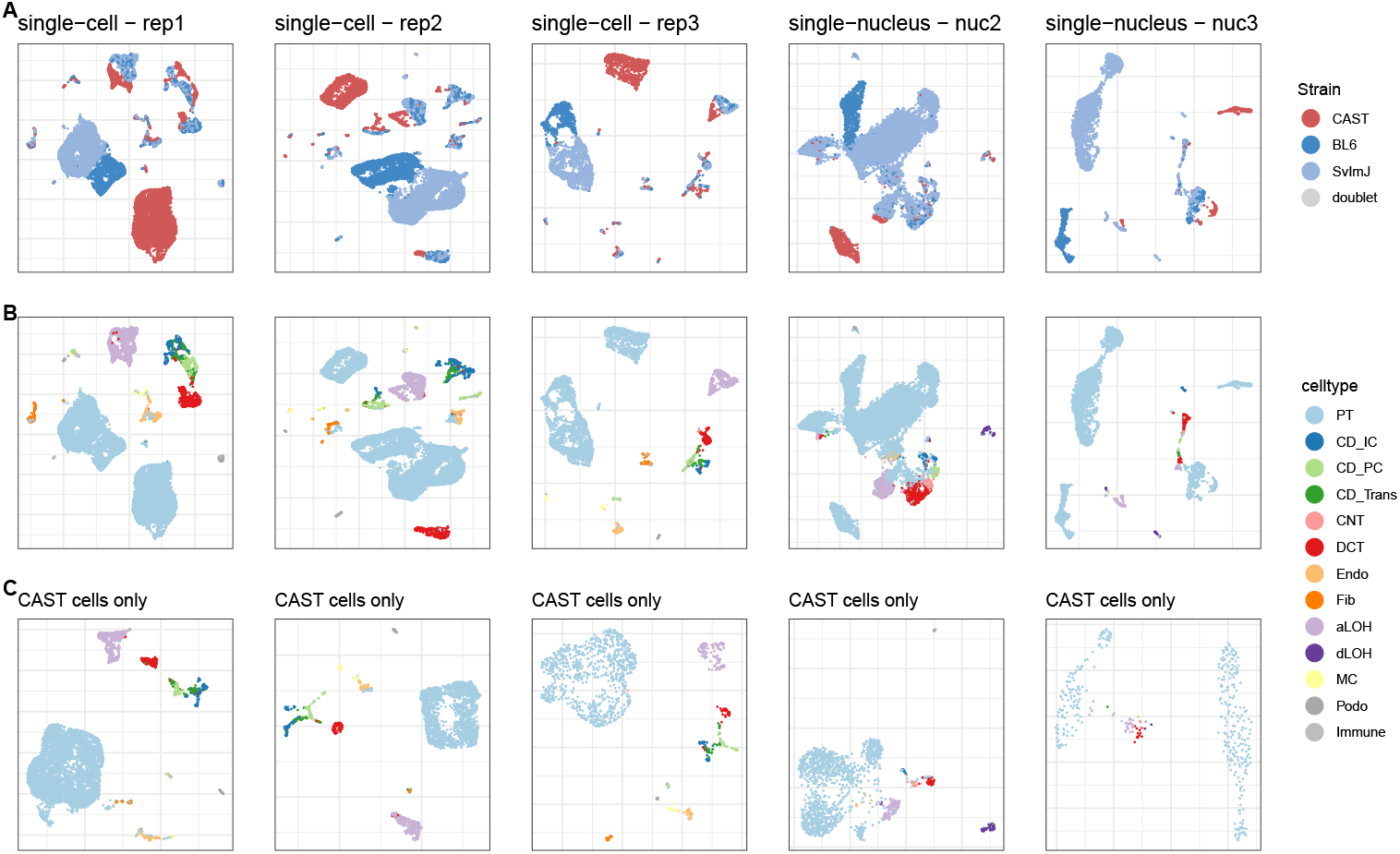
UMAP visualization showing the composition per replicate of. A) all cells, colored by strain assignment, B) all cells, colored by cell type assignment and C) *M. m. castaneus* cells only, colored by cell type assignment.

**Supplementary Figure S3.**
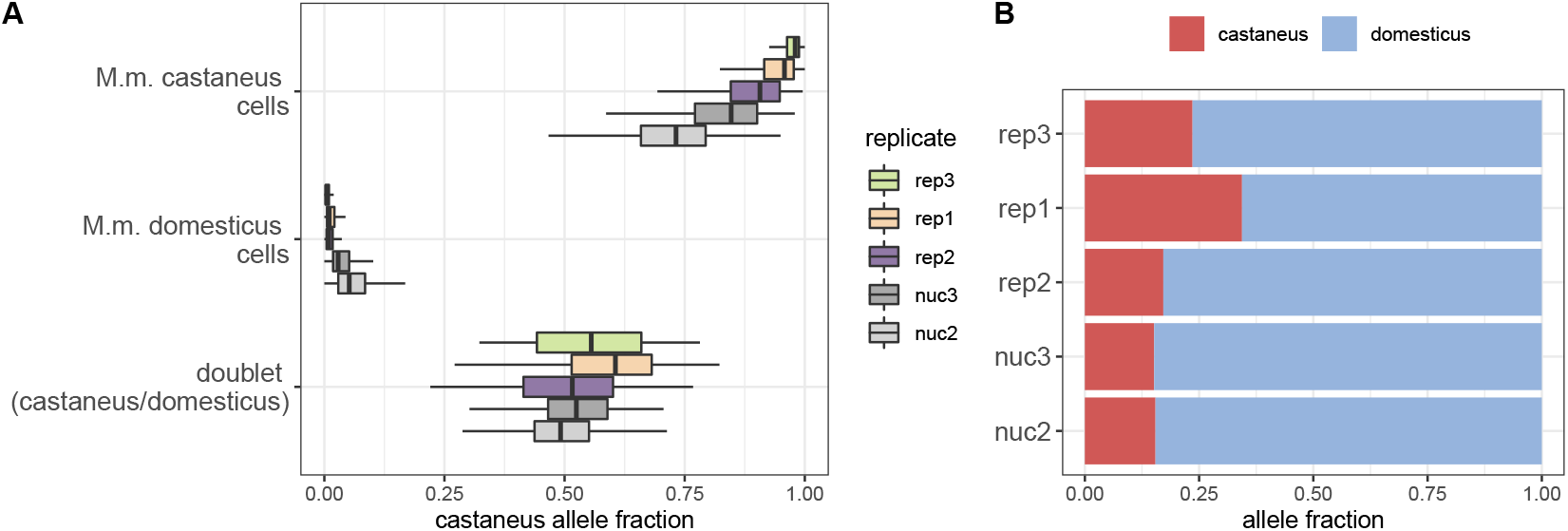
Detection of cross-genotype contamination. A) *M*.*m. castaneus* allele frequency per cell in cells from different subspecies and mixed-subspecies doublets. In all replicates varying amounts of *M*.*m. castaneus* alleles are detected in *M*.*m. domesticus* cells and vice versa, pointing towards background noise originating from cross-genotype contamination. B) Allele frequency proportions across all cells in a replicate.

**Supplementary Figure S4.**
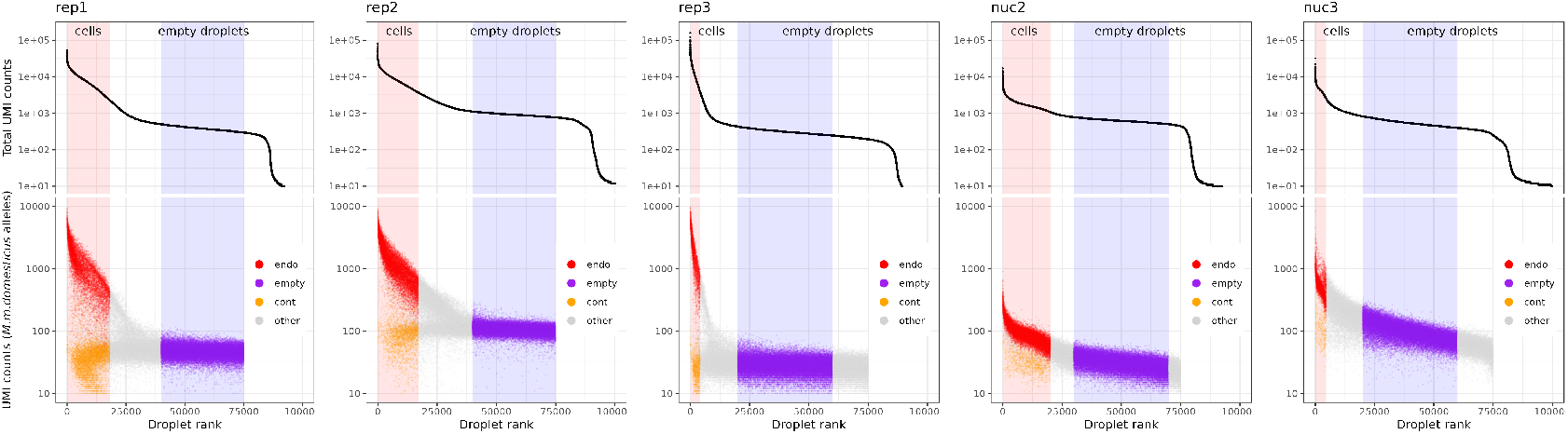
Definition of endogenous, empty droplet and contamination profiles across replicates. Droplet barcodes were ordered by their total UMI counts and empty droplets were defined from this UMI curve as barcodes in the low UMI count plateau area (upper panel). UMI counts of reads covering *M. m. domesticus* specific alleles were used to construct three different profiles (lower panel). *M. m. domesticus* allele counts in *M. m. domesticus* cells were defined as endogenous counts (endo), *M. m. domesticus* allele counts in *M. m. castaneus* cells as contaminating counts (cont) and *M. m. domesticus* allele counts associated with barcodes of the empty droplet plateau as empty droplet counts (empty).

**Supplementary Figure S5.**
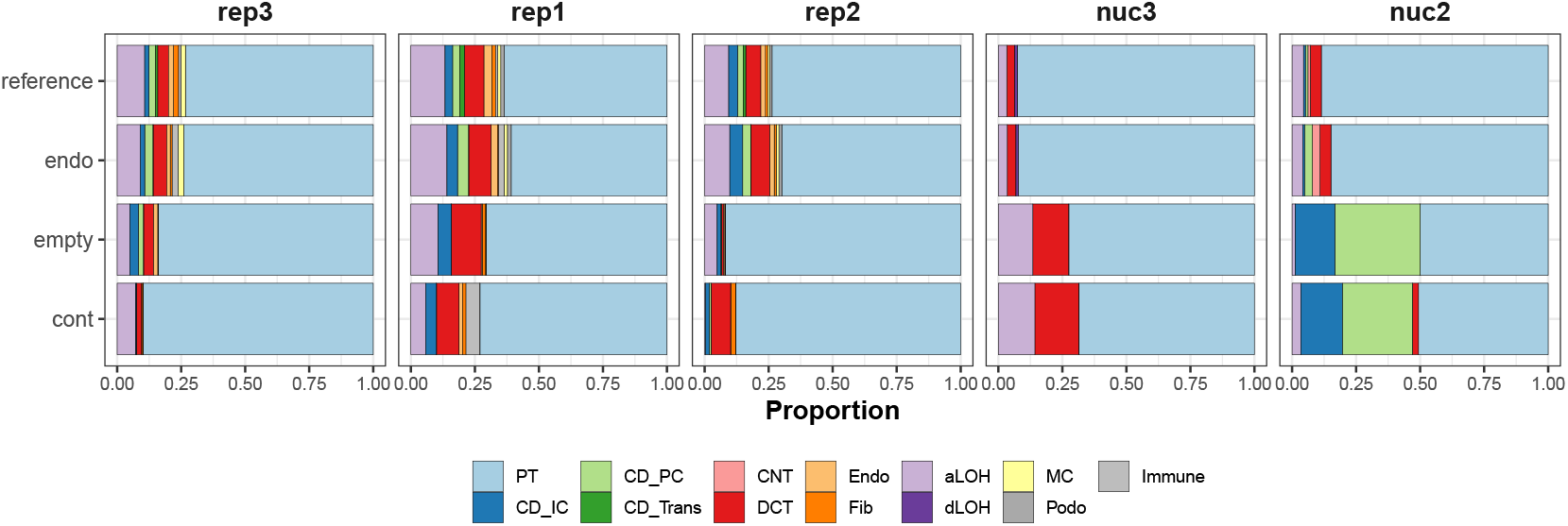
Dissection of cell type contributions by deconvolution of pseudobulk profiles. The stacked bar plots of ‘reference’ depict the proportions of cell types in a single cell reference used for deconvolution with SCDC [16]. The ‘endo’,’empty’ and ‘cont’ bar plots show the estimated fraction of cell types after deconvolution of pseudobulk profiles that were aggregated for each category.

**Supplementary Figure S6.**
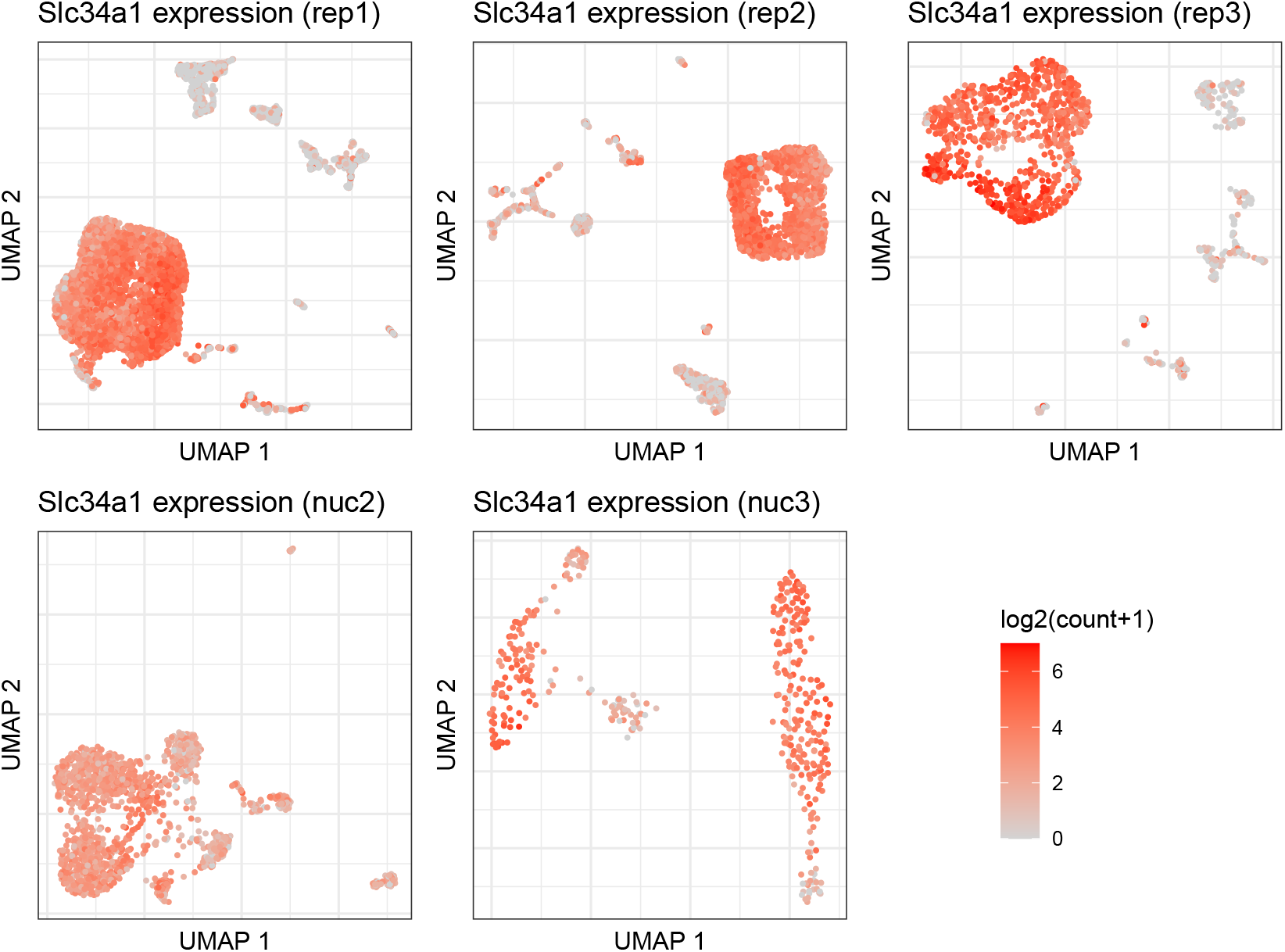
Slc34a1 expression across replicates. UMAP representation *M. m. castaneus* cells coloured by Slc34a1 expression. Spurious detection of Slc34a1 in all cell clusters is observed in all replicates. In the replicates with the lowest background noise levels (rep1,rep3), Slc34a1 expression is most concentrated in PT cells.

**Supplementary Figure S7.**
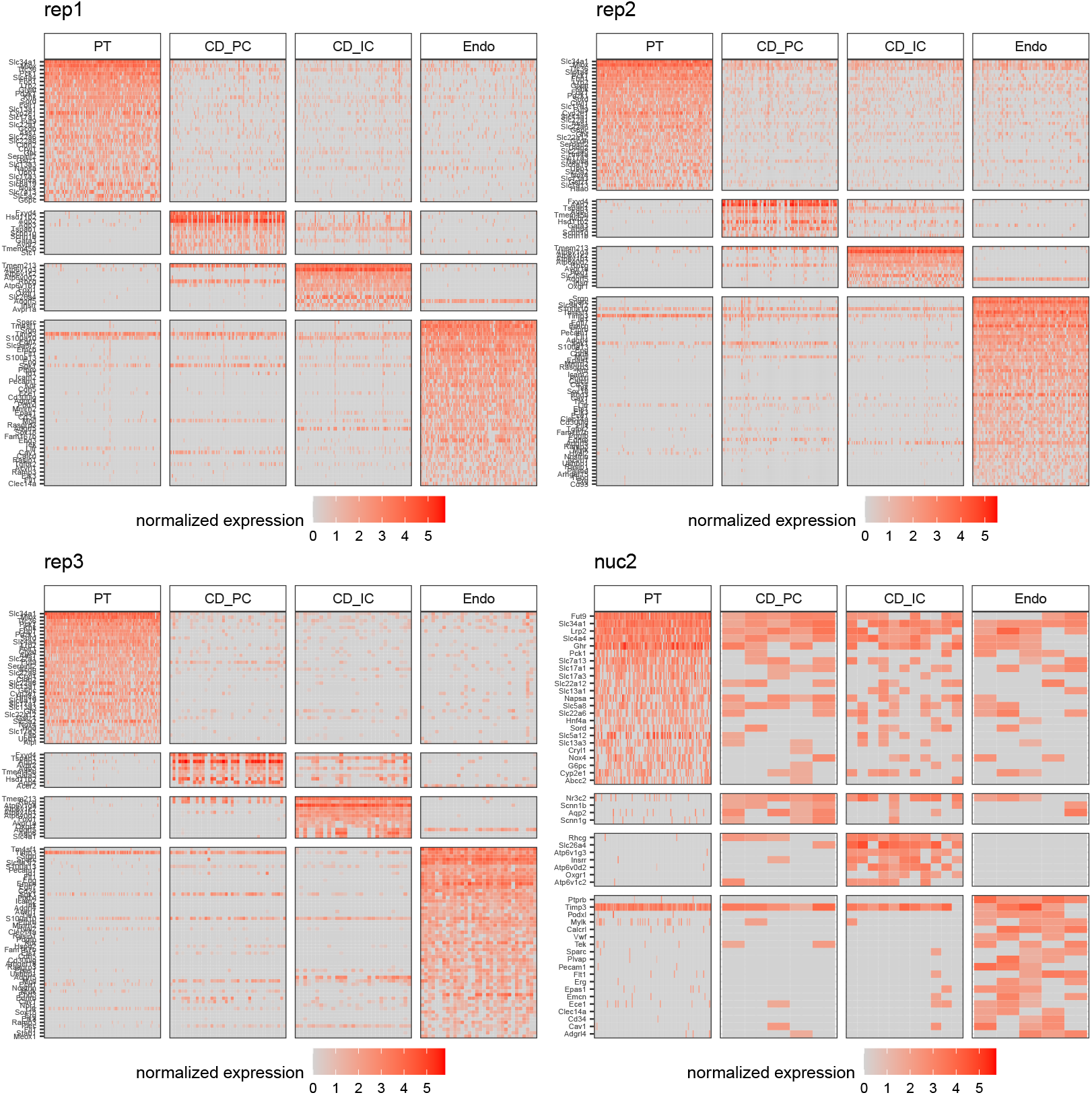
Expression of cell type marker genes. Heatmaps show the normalized expression of known marker genes for four selected cell types across replicates. Marker genes were obtained from the PanlaoDB database [17] and filtered to select for genes that are detected in at least 50% of the cells of the cell type in which they are expected to be expressed. The replicate nuc3 was excluded from this figure due to an insufficient number of collecting duct and endothelial cells. PT: proximal tubule; CD_IC: intercalated cells of collecting duct; CD_PC: principal cells of collecting duct; Endo: endothelial

**Supplementary Figure S8.**
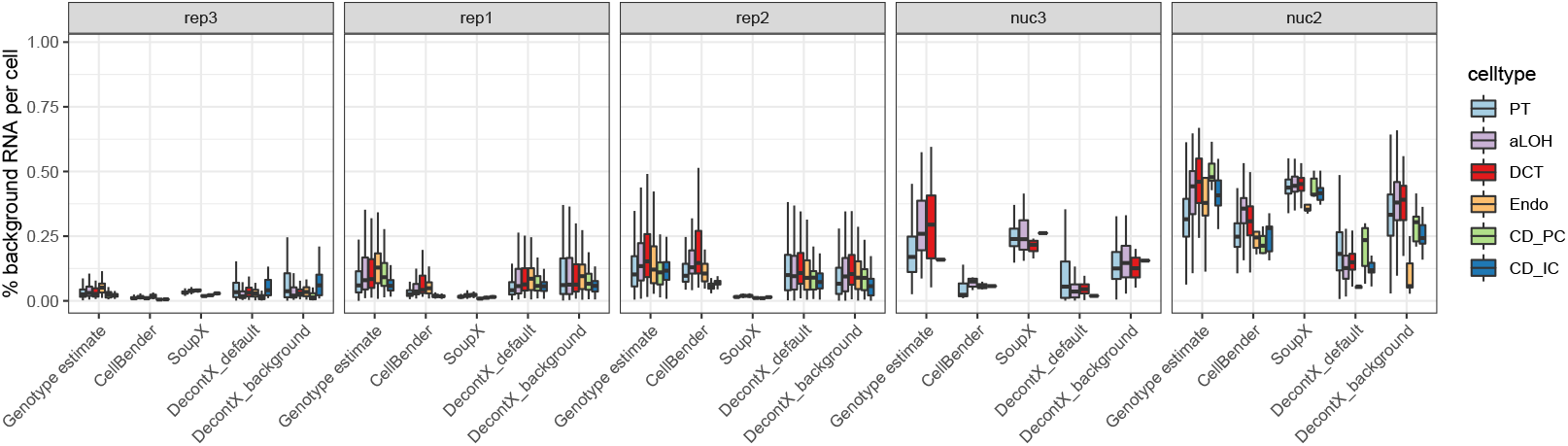
Estimated background noise levels across cell types. Genotype estimates are inferred based on genetic variants. Cellbender, SoupX and DecontX estimates are calculated for each cell based on a corrected count matrix. PT: proximal tubule; aLOH: ascending loop of Henle; DCT: distal convoluted tubule; Endo: endothelial; CD_PC: principal cells of collecting duct; CD_IC: intercalated cells of collecting duct.

**Supplementary Figure S9.**
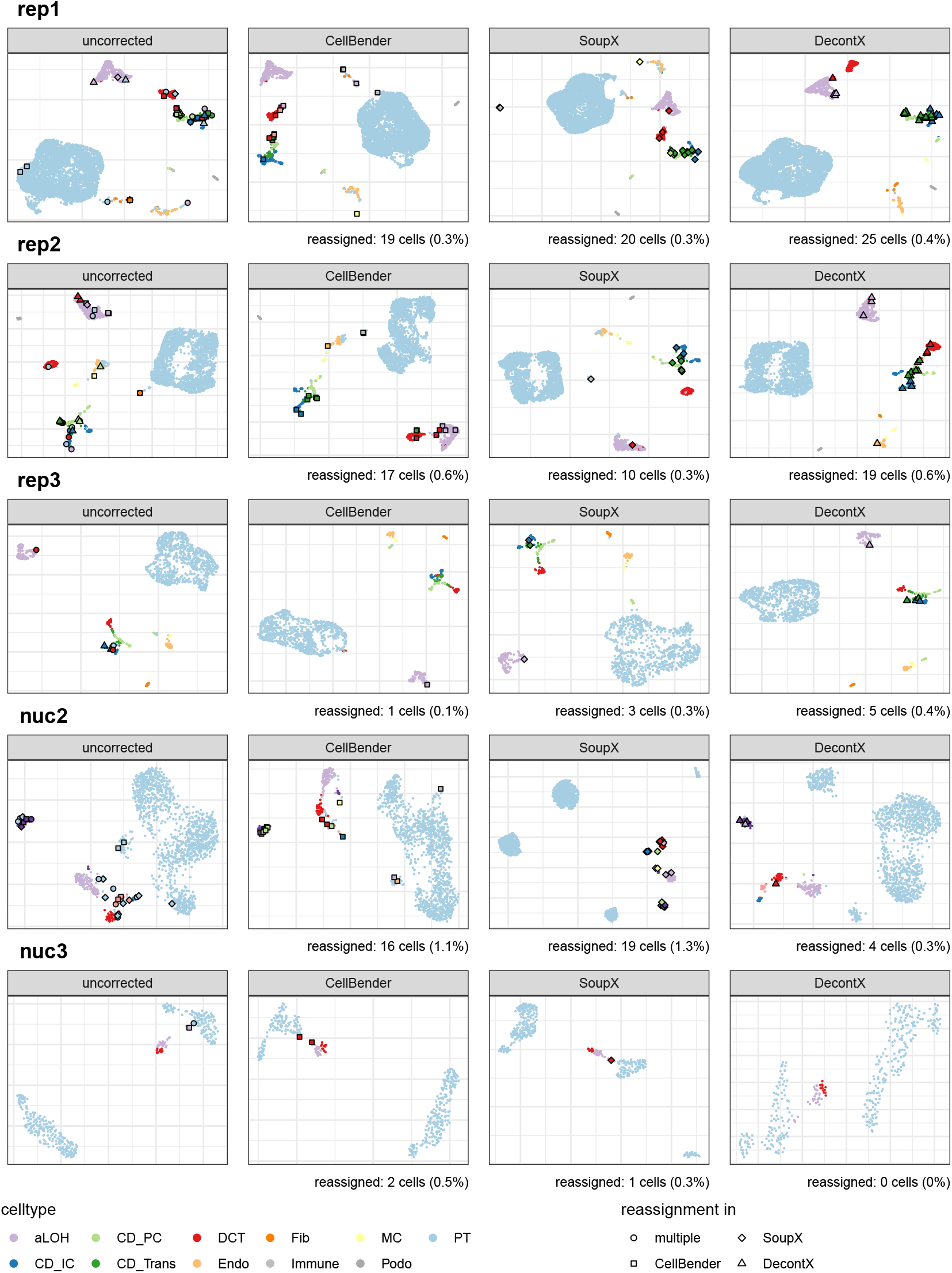
UMAP representations of all replicates before and after background noise correction. Cells are colored by cell type labels obtained from reference based classification. Individual cells that received a new label after correction are highlighted. In case of the uncorrected data, all cells that received a new label after correction with any of the methods are highlighted. PT: proximal tubule; CD_IC: intercalated cells of collecting duct; CD_PC: principal cells of collecting duct; CD_Trans: transitional cells of collecting duct; CNT: connecting tubule; DCT: distal convoluted tubule; Endo: endothelial; Fib: fibroblasts; aLOH: ascending loop of Henle; dLOH: descending loop of Henle; MC: mesangial cells; Podo: podocytes

**Supplementary Figure S10.**
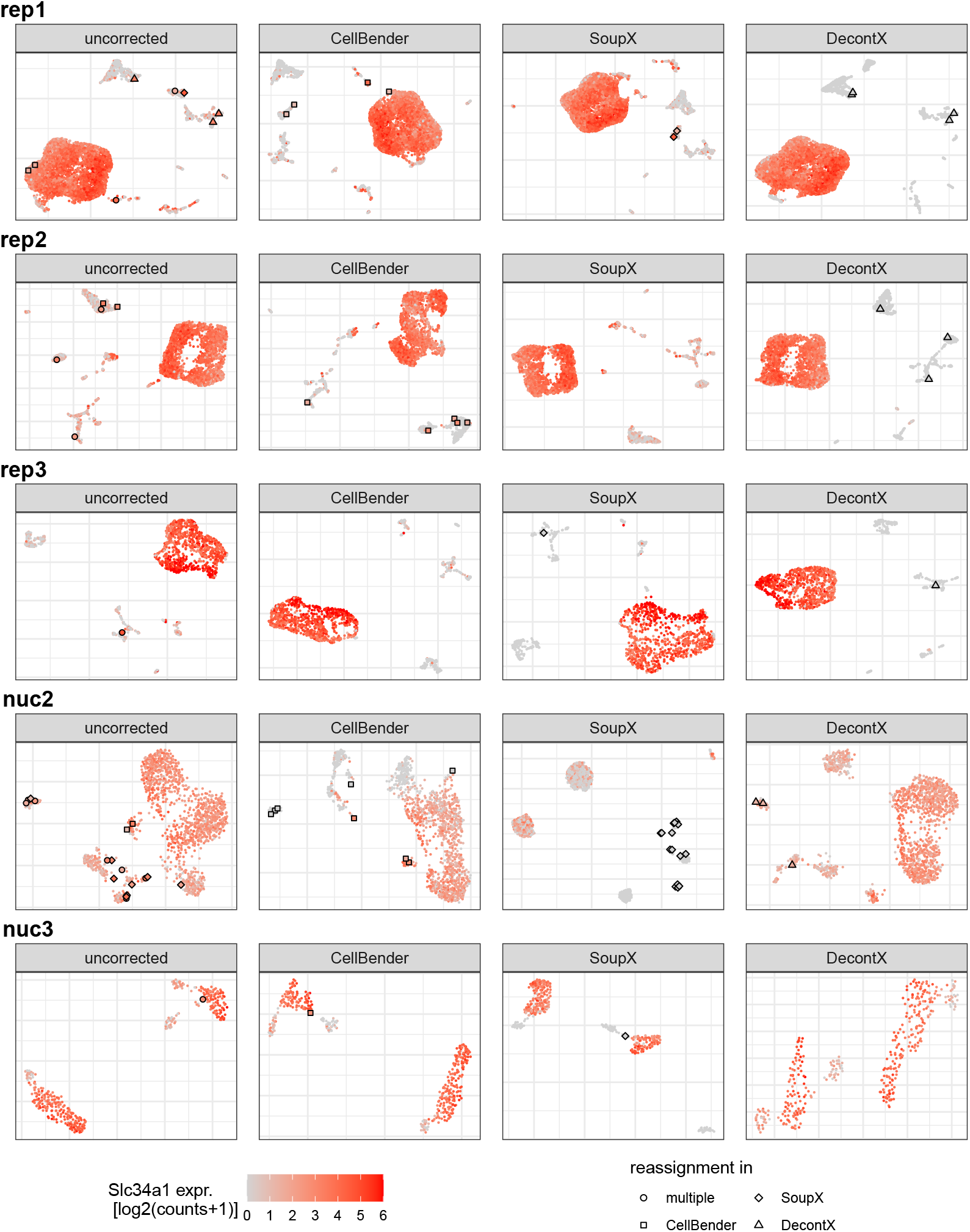
Detected expression levels of Slc34a1 before and after background noise correction. Cells that were classified as PT cells in the uncorrected data, but got reassigned after correction, are highlighted.

**Supplementary Figure S11.**
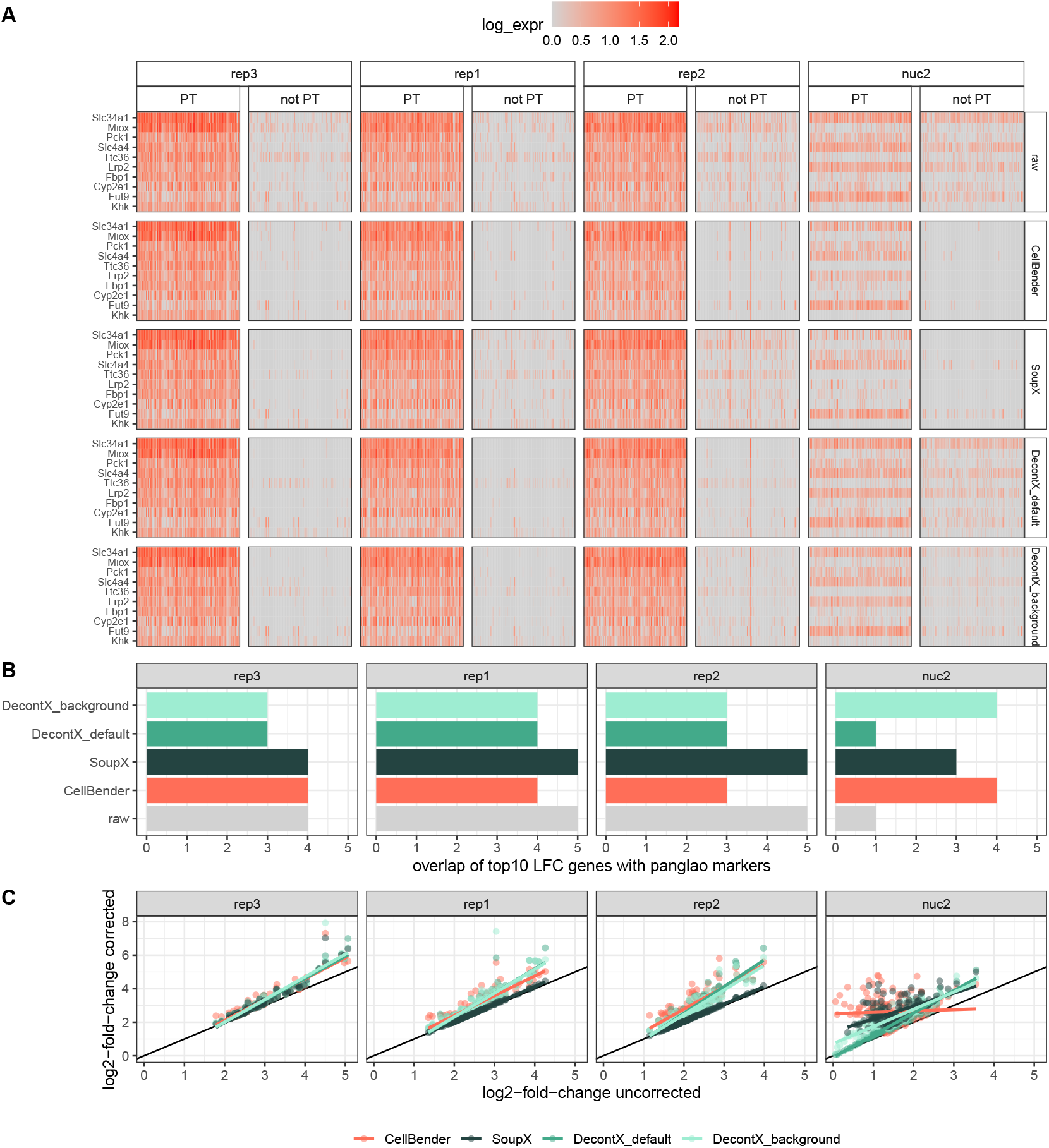
Effect of background noise correction on marker gene detection. A) Heatmaps depicting the expression of 10 PT marker genes in 100 randomly sampled PT cells and 100 cells from other cell types. The first row of heatmaps is based on the uncorrected count matrix, rows 2-5 on the denoised count matrix output by different methods. B) Overlap of identified and known marker genes. Genes were ranked by log2 fold change between PT an other cells and the overlap of the top 10 genes in this ranking with known marker genes for Proximal Tubule cells from PanglaoDB [17] is shown. C) Log2 fold changes of PangloaDB PT cell marker genes after background noise correction compared to the uncorrected data.

**Supplementary Figure S12.**
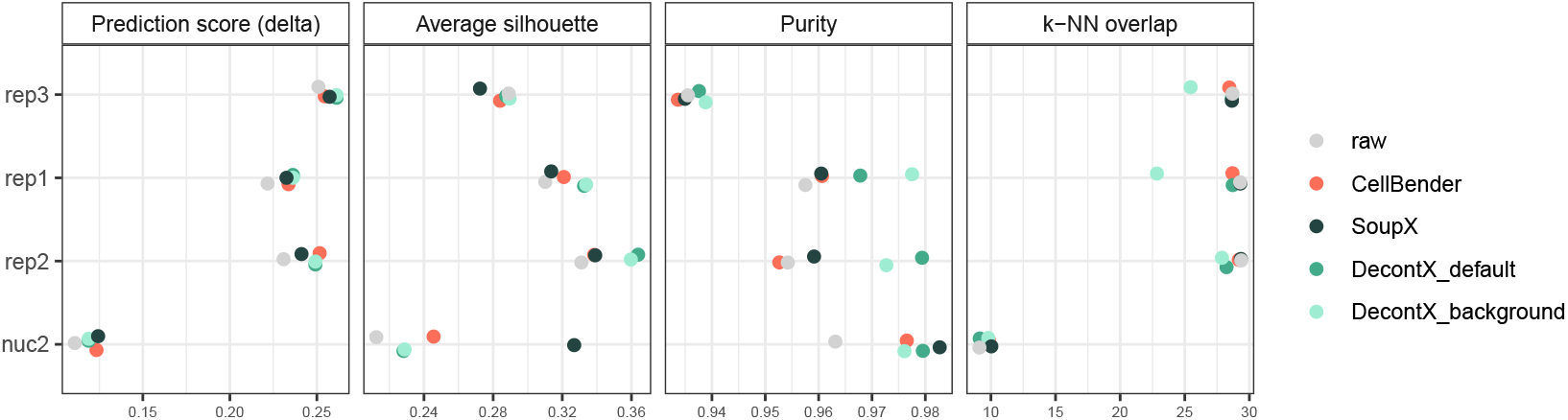
Evaluation metrics for cell type identification. Prediction score: cell-wise score “delta” of reference based classification with SingleR [21]. Average silhouette: Mean of silhouette widths per cell type. Purity: Cluster purity calculated on cell type lables as ground truth and Louvain clusters as test labels. *k* -NN overlap: overlap of the *k* =50 nearest neighbors per cell compared to genotype-cleaned reference *k* -NN graph.

**Supplementary Figure S13.**
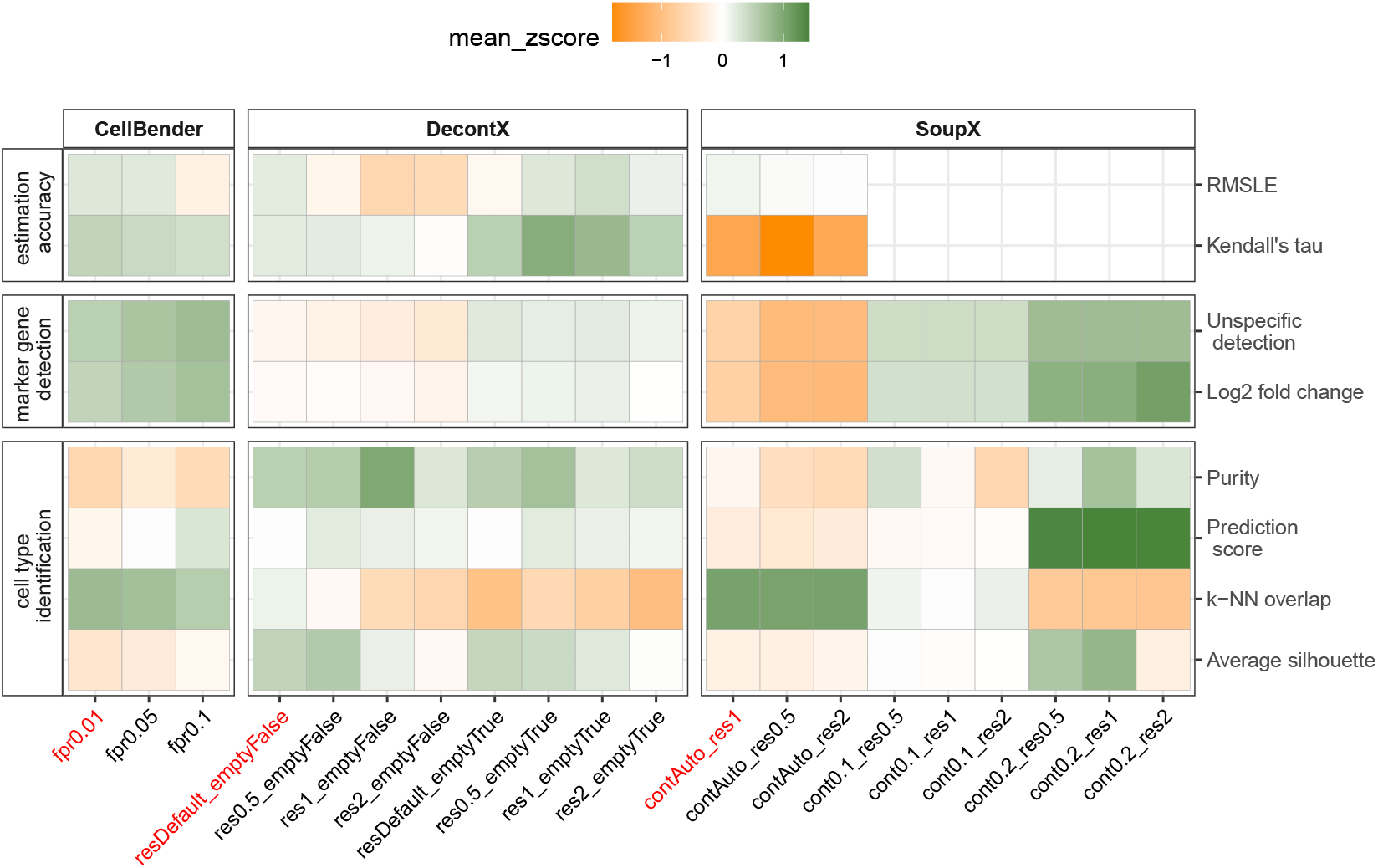
Evaluation of different parameter settings. Combinations of the most impactful parameter/workflow choices of each method are evaluated. Default parameter settings are highlighted with red font color. For each metric, an average z-score across the replicates rep1, rep2, rep3 and nuc2 is shown, for which higher values indicate better performance. The following parameters were tuned: CellBender: *fpr* (0.01,0.05,0.1); DecontX: cluster lables *z* (resDefault: NULL, res0.5/1/2: vector of cluster labels from Louvain clustering with resolution 0.5/1/2), *background* (emptyFalse: NULL, emptyTrue: provide raw matrix containing empty droplets); SoupX: contamination fraction (contAuto: automatic estimation using *autoEstcont*, cont0.1/0.2: manually set using *setContaminationFraction* (0.1/0.2)), cluster labels (res0.5/1/2: vector of cluster labels from Louvain clustering with resolution 0.5/1/2)

